# Substitutions at Non-Conserved Rheostat Positions Modulate Function by Re-Wiring Long-Range, Dynamic Interactions

**DOI:** 10.1101/2020.07.02.185207

**Authors:** Paul Campitelli, Liskin Swint-Kruse, S. Banu Ozkan

## Abstract

Amino acid substitutions at nonconserved protein positions can have non-canonical and “long-distance” outcomes on protein function. Such outcomes might arise from changes in the internal protein communication network, which is often accompanied by changes in structural flexibility. To test this, we calculated flexibilities (“DFI”) and dynamic coupling (“DCI”) for positions in the linker region of the lactose repressor protein (“LacI”). This region contains nonconserved positions for which substitutions alter DNA binding affinity. We first chose to study eleven substitutions at position 52. In computations, substitutions showed long-range effects on flexibilities of DNA binding positions, and the degree of flexibility change correlated with experimentally-measured changes in DNA binding. Substitutions also altered dynamic coupling to DNA binding positions in a manner that captured other experimentally-determined functional changes. Next, we broadened calculations to consider the dynamic coupling between 17 linker positions and the DNA binding domain. Experimentally, these linker positions exhibited a wide range of substitution outcomes: Four conserved positions tolerated almost no substitutions (“toggle”), ten nonconserved positions showed progressive changes from a range of substitutions (“rheostat”), and three nonconserved positions tolerated almost all substitutions (“neutral”). In computations with wild-type LacI, the dynamic couplings between the DNA binding domain and these linker positions showed varied degrees of asymmetry that correlated with the observed toggle/rheostat/neutral substitution outcomes. Thus, we propose that long-range and non-canonical substitutions outcomes at nonconserved positions arise from re-wiring long-range communication among functionally-important positions. Such calculations might enable predictions for substitution outcomes at a range of nonconserved positions.

## Introduction

Historically, protein substitution experiments have been biased towards amino acid positions that are conserved during evolution (Kumar et al. 2009). Changes at these positions are often disease-causing and much effort has been spent towards understanding and predicting their outcomes. However, advancements in genome sequencing have also shown thousands of variations at nonconserved protein positions. By analogy to paralog evolution, changes at these positions can also alter protein function. Predicting the outcome of mutations at nonconserved positions, particularly for cases that do not cause large-scale structural differences from the wild-type structure, remains a considerable challenge (Miller et al. 2017; Swint-Kruse 2016).

We previously used the lactose repressor protein (“LacI”, fig. 1) as a model system for understanding the functional roles of nonconserved positions. In this work, we focused on nonconserved positions in the linker region that forms interfaces with the two major LacI functional domains (Bell and Lewis 2000; Swint-Kruse et al. 2002). Many linker positions are nonconserved (supplementary table S1) and their substitutions often showed non-canonical outcomes (Meinhardt et al. 2013): When a range of amino acids were substituted into each of the positions, the resulting changes in transcription repression and/or DNA binding affinity ranged several orders of magnitude (*e*.*g*. supplementary fig S1; table 1). Notably, when the functional data for each position’s substitutions were arranged by their rank order, the resulting plots did not correlate with either side chain chemistries nor could be explained by side chain characteristics (including their effects on helical propensity, hydrophobicity, side chain size, etc.) or evolutionary frequency (Meinhardt et al. 2013; Zhan et al. 2006). Since each variant was capable of binding DNA (Meinhardt et al. 2013), we reasoned that structural rearrangements must be minimal. However, previous experimental and computational studies indicated that the LacI linker is very flexible and that changes in its folding (and thus flexibility) are integral to LacI function (Ha et al. 1989; Kalodimos et al. 2001; Kalodimos et al. 2004; Kalodimos, Bonvin et al. 2002; Lewis et al. 1996; Spolar and Record 1994; Spronk et al. 1996; Swint-Kruse et al. 2002; Taraban et al. 2008; Wade-Jardetzky et al. 1979). Thus, we hypothesized that altered dynamics contribute the non-canonical outcomes that arise from amino acid variation in the LacI linker.

**Table 1.**
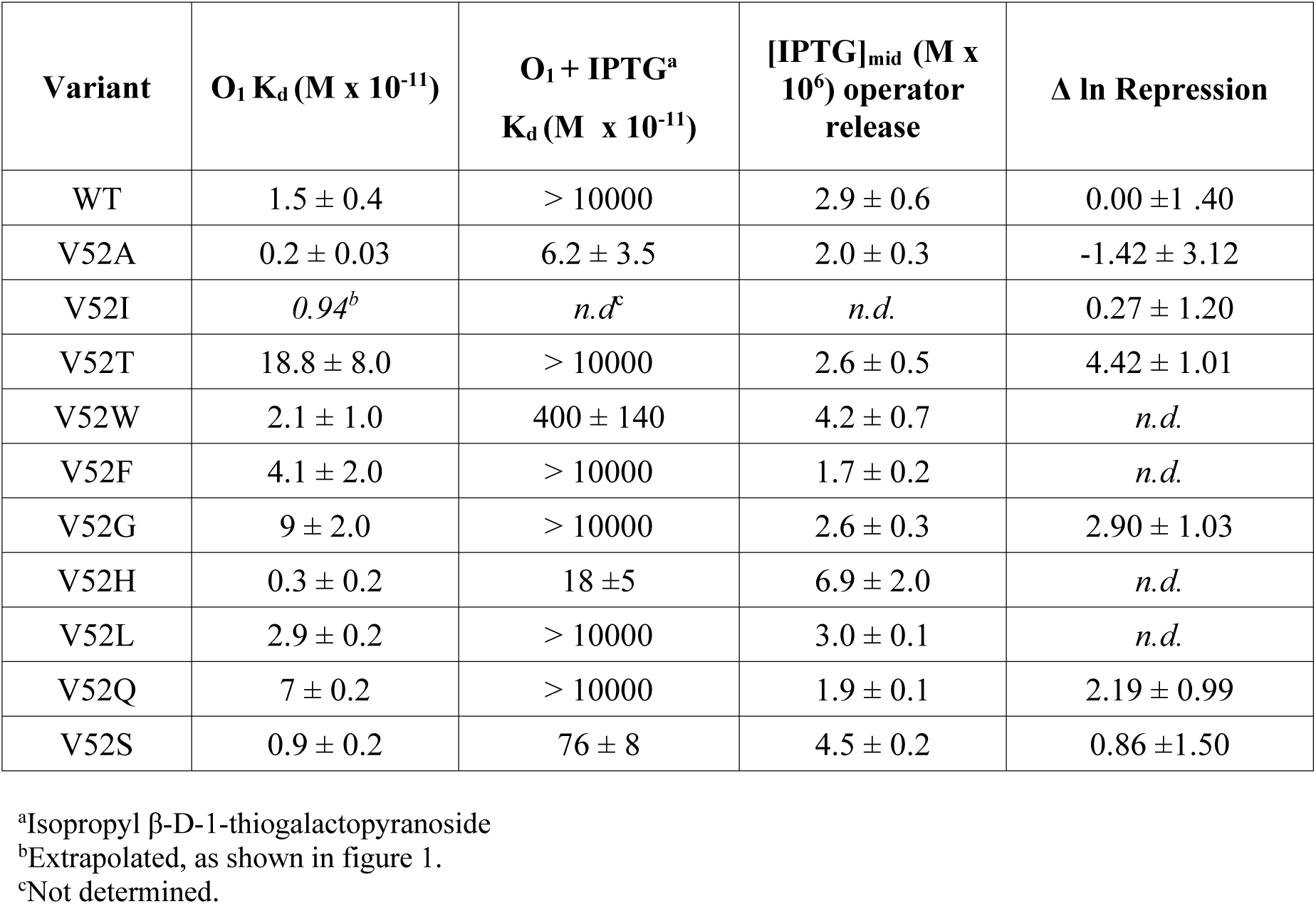
Experimentally measured functional parameters for LacI V52X variants. (Meinhardt et al. 2013; Zhan et al. 2006).

| Variant | $O_1 K_d (M \times 10^{-11})$ | $O_1 + IPTG^a$<br>$K_d (M \times 10^{-11})$ | $[IPTG]_{mid} (M \times 10^6)$ operator release | $\Delta \ln Repression$ |
| --- | --- | --- | --- | --- |
| WT | $1.5 \pm 0.4$ | $> 10000$ | $2.9 \pm 0.6$ | $0.00 \pm 1.40$ |
| V52A | $0.2 \pm 0.03$ | $6.2 \pm 3.5$ | $2.0 \pm 0.3$ | $-1.42 \pm 3.12$ |
| V52I | $0.94^b$ | $n.d.^c$ | $n.d.$ | $0.27 \pm 1.20$ |
| V52T | $18.8 \pm 8.0$ | $> 10000$ | $2.6 \pm 0.5$ | $4.42 \pm 1.01$ |
| V52W | $2.1 \pm 1.0$ | $400 \pm 140$ | $4.2 \pm 0.7$ | $n.d.$ |
| V52F | $4.1 \pm 2.0$ | $> 10000$ | $1.7 \pm 0.2$ | $n.d.$ |
| V52G | $9 \pm 2.0$ | $> 10000$ | $2.6 \pm 0.3$ | $2.90 \pm 1.03$ |
| V52H | $0.3 \pm 0.2$ | $18 \pm 5$ | $6.9 \pm 2.0$ | $n.d.$ |
| V52L | $2.9 \pm 0.2$ | $> 10000$ | $3.0 \pm 0.1$ | $n.d.$ |
| V52Q | $7 \pm 0.2$ | $> 10000$ | $1.9 \pm 0.1$ | $2.19 \pm 0.99$ |
| V52S | $0.9 \pm 0.2$ | $76 \pm 8$ | $4.5 \pm 0.2$ | $0.86 \pm 1.50$ |

<sup>a</sup>Isopropyl $\beta$ -D-1-thiogalactopyranoside
<sup>b</sup>Extrapolated, as shown in figure 1.
<sup>c</sup>Not determined.

**Figure 1.**
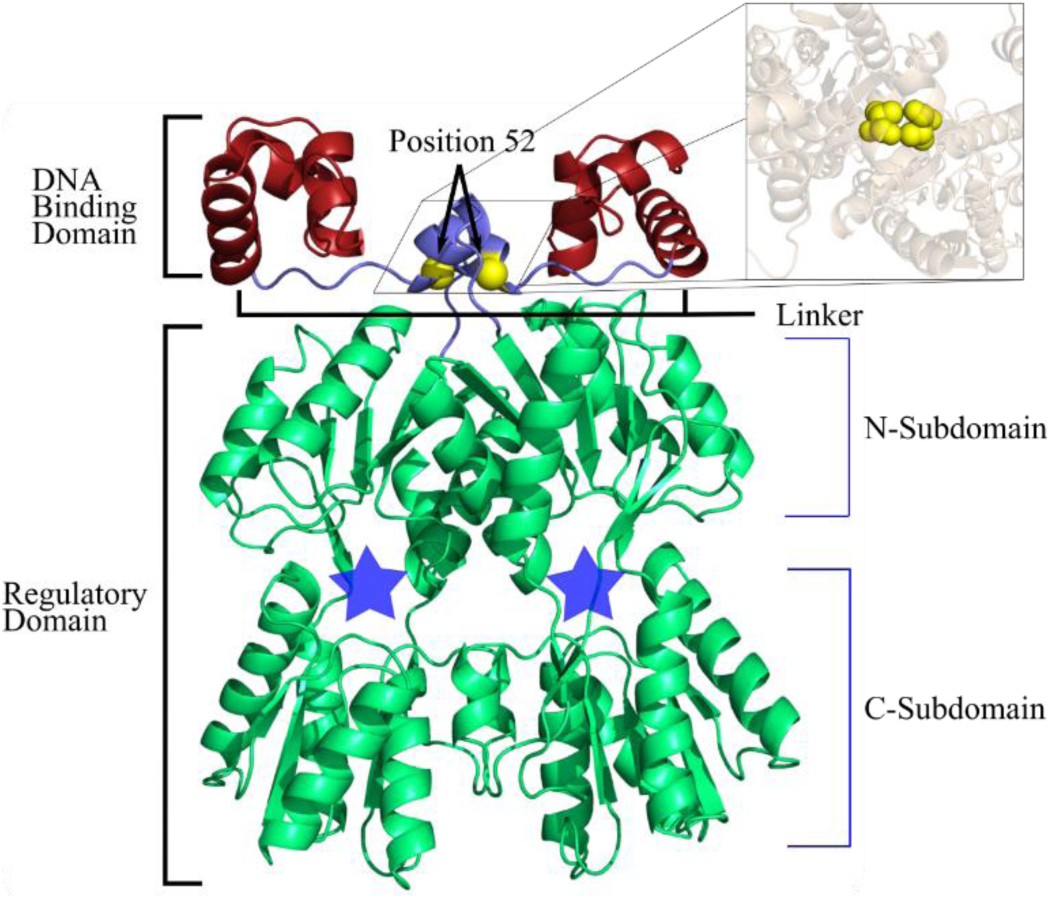
LacI structure (pdb ID 1efa; (Bell and Lewis 2000)) colored and labeled by domain. The minimal LacI structure for high affinity DNA binding is a homodimer (reviewed in (Swint-Kruse and Matthews 2009)). Each monomer comprises two major domains – a DNA binding domain (red) and a regulatory domain (green) – that are connected by a flexible linker region (blue). DNA binding involves positions on the top surface of the DNA binding domains as well as several linker positions. The Cα’s of linker position 52 are highlighted with yellow spheres; the inset shows a top-down view of both V52 side chains using space-filling spheres. In the regulatory domains, locations of the allosteric effector binding sites are indicated with blue stars. Small molecule “inducer” binding to this site diminishes DNA binding affinities by up to 1000-fold (Swint-Kruse and Matthews 2009). The structure shown and used in this work was extracted from a crystal structure (1efa, (Bell and Lewis 2000)) that was determined in the presence of bound DNA and ortho-nitrophenyl-beta-d-fucopyranoside (“ONPF”, an anti-inducer that binds in the allosteric site and enhances DNA binding). This figure was rendered with pyMOL (Schrodinger LLC 2010).

To test this hypothesis, we performed in-depth analyses for substitutions at LacI linker position 52. We chose this position because, among all linker positions, 52 had the widest range of substitution outcomes and the most extensive biochemical data available (Hodges et al. 2018; Sousa et al. 2016; Zhan et al. 2006). Experimentally, substitution outcomes for this position spanned more than 4 orders of magnitude (Meinhardt et al. 2013; Zhan et al. 2006) (*e*.*g*. table 1).

*In vitro* DNA binding affinities (K_d_ values) were available for wild-type LacI and 11 variants, of which six overlapped with the set of variants characterized by quantitative *in vivo* repression assays. Repression showed a strong correlation with K_d_ (supplementary fig. S1), which suggests that repression changes were due to altered DNA binding affinity. From these two experimental datasets, we used eleven variants (including wild-type LacI), which we designate as the “V52X” set throughout this text.

Here, we explored whether V52X substitutions affected flexibility at mechanistically important regions of the LacI structure and/or altered dynamic coupling between the DNA binding domain and the linker. To that end, we first calculated site-specific flexibility of wild-type LacI and V52X variants using the dynamic flexibility index (“DFI”). The DFI parameter measures each position’s sensitivity to perturbations within a network of interactions and represents a given amino acid position’s ability to explore its local conformational space (flexibility). For LacI, the flexibilities of the DNA binding domain and linker region differed significantly among the V52X variants, and the flexibility changes correlated strongly with changes in DNA binding affinities.

Linker position 52 does not directly contact DNA and, indeed, is distal to many DNA binding domain positions.Thus, we further reasoned that the flexibility changes observed in the DNA binding domain must be propagated from the V52X variants via altered long-range dynamic coupling. To that end, we calculated the dynamic coupling index (“DCI”) for these variants. DCI measures the displacement response of an individual position to the perturbation of a second position or group of positions, relative to the average response to any perturbation of all possible positions. For each of the V52X variants, coupling to positions in the DNA binding domain did indeed show different profiles, suggesting that changes in long-range dynamic coupling may underlie the altered flexibilities of the DNA binding domains.

Given the compelling results for this representative nonconserved position, we next compared DFI and DCI calculations for all 13 nonconserved positions in the wild-type LacI linker to four conserved linker positions. Within the linker region (positions 46-62), 13 positions were varied among homologs in the broader LacI/GalR family (Tungtur et al. 2011) and exhibited a range of sequence entropies (Parente and Swint-Kruse 2013). In contrast, the other four linker positions were very strongly conserved across the whole family (Parente and Swint-Kruse 2013; Tungtur et al. 2011) with low sequence entropies (supplementary table S1). Most of the nonconserved positions had substitution outcomes similar to position 52, with intermediate changes in biochemical function (Meinhardt et al. 2013), but the range of change varied from <1 to >4 orders of magnitude. In contrast, the four conserved linker positions showed canonical substitution outcomes, with most substitutions abolishing repression (Markiewicz et al. 1994; Miller et al. 2017; Suckow et al. 1996). Neither DFI nor DCI values discriminated these patterns of substitution outcomes. However, DCI values measure coupling that is inherently asymmetric: Every amino acid position has a unique network of direct, local interactions that gives rise to a unique network of highly coupled partner positions (fig. 2). Across the protein structure, this gives rise to an inhomogeneous, overall 3-D network, and the asymmetry provides directionality to long-distance coupling. Thus, for a particular pair of coupled amino acids (*i* and *j*), their unique network constraints differentiate the coupling of *i* to *j* from the coupling of *j* to i. When DCI analyses were used to assess asymmetric coupling between linker positions and those in the DNA binding domain, distinct patterns were identified that correlated with different patterns of substitution outcomes.

**Figure 2.**
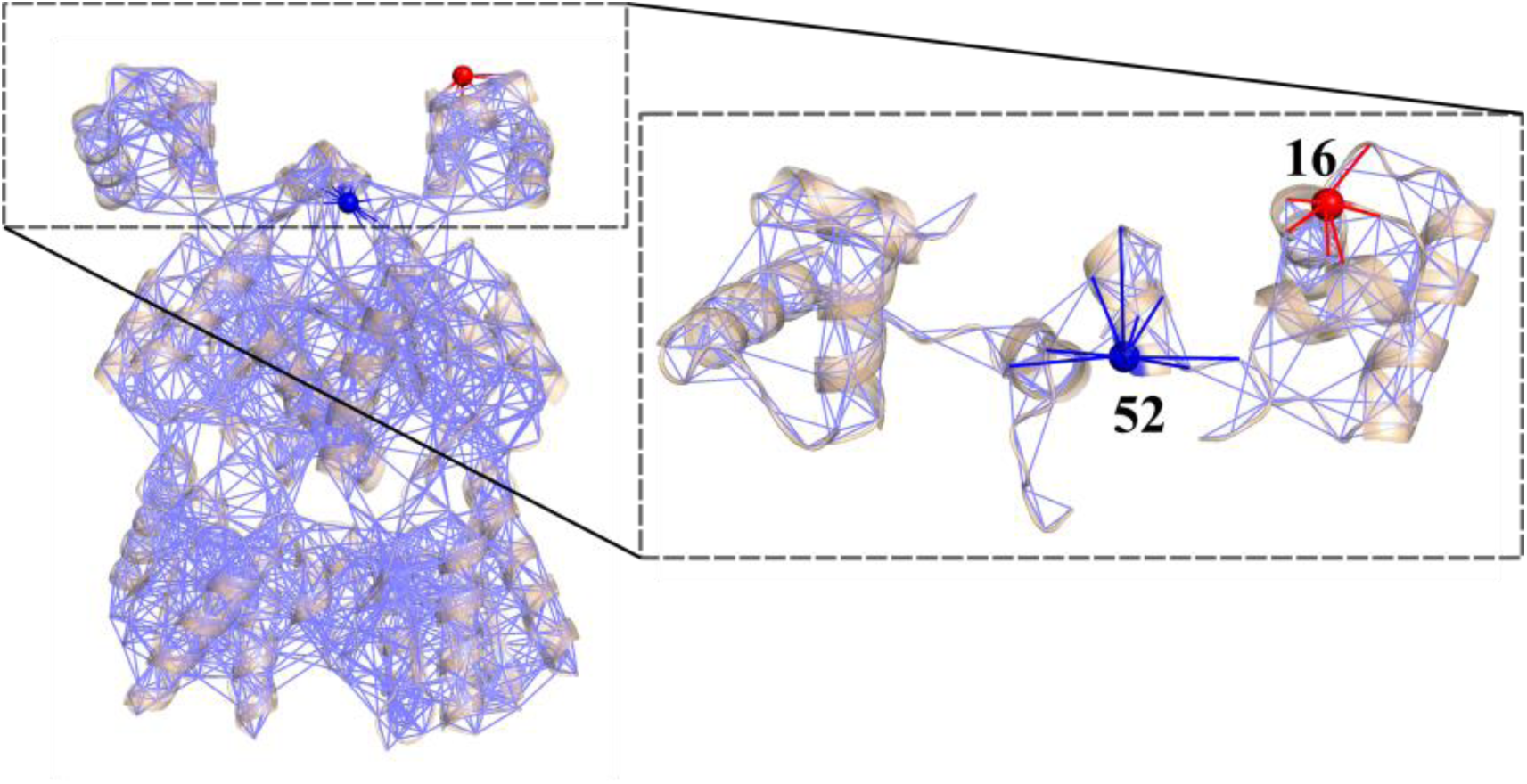
Protein structure interactions can be represented as inhomogeneous networks. In this example using LacI, each amino acid is represented as a node in the network, and covalent and noncovalent interactions are represented as edges. The expanded area in the dashed box shows a close-up view of the DNA binding domains and linkers, with two positions highlighted (blue and red) to exemplify asymmetric coupling. Due to the inhomogenous interaction networks between positions 52 and 16 and their partner positions, a perturbation at the red site may induce a response at the blue site that differs in magnitude from the response arising at the red site with the blue site is perturbed.

Taken together, the current analyses provide a rationale for the substitution outcomes at nonconserved linker positions for which no previous relationships could reconcile and suggest that these positions are nodes in the dynamic communication network that exists between the functional domains of LacI.

## Results

The dynamic flexibility index (DFI) combines Perturbation Response Scanning and Linear Response Theory to evaluate each position’s displacement response to random force perturbations at other locations in the protein (Gerek and Ozkan 2011; Nevin Gerek et al. 2013). For each position *i*, the DFI value quantifies its total displacement response relative to the total displacement response for all positions in the protein. To identify the most flexible positions, ranked *%*DFI values are used: For each position *i*, its percentile rank in the observed DFI range is calculated from the ratio of the number of positions with DFI values ≤ DFI_i_ to the total number of positions. *%*DFI values range from 0 to 1. The DFI parameter can be considered a measure of a given position’s ability to explore its local conformational space; that is, DFI is a measure of an amino acid’s flexibility. This parameter has been used to identify important structural elements of a protein, such as hinges, and to show the flexibility of functionally critical areas, such as binding domains and catalytic regions (Bolia et al. 2012; Bolia and Ozkan 2016; Campitelli et al. 2018; Gerek and Ozkan 2010, 2011; Kumar et al. 2015; Modi et al. 2018; Modi and Ozkan 2018; Nevin Gerek et al. 2013).

Figure 3 presents the *%*DFI profile for wild-type LacI mapped onto its structure, using a color spectrum of red (highest flexibility) to blue (lowest flexibility). Calculations were performed using the protein structure extracted from the DNA-bound crystal structure 1efa (Bell and Lewis 2000) (*i*.*e*. no DNA was present in the calculation). The calculated DFI profile indicates that the wild-type structure exhibits a relatively flexible linker region and DNA binding domain and a comparably stiff regulatory domain. The two subunits of the homodimer show symmetry in their flexibilities. These results agree with a variety of structural and biochemical evidence: As mentioned in the Introduction, a range of studies (Ha et al. 1989; Kalodimos et al. 2001; Kalodimos et al. 2004; Kalodimos, Bonvin et al. 2002; Lewis et al. 1996; Spolar and Record 1994; Spronk et al. 1996; Swint-Kruse et al. 2002; Taraban et al. 2008; Wade-Jardetzky et al. 1979) indicate flexibility of the DNA binding domains and linker regions. In addition, comparisons of crystal structures for different ligand-bound forms of LacI show that the C-subdomains of the regulatory domains remain fixed, whereas the two N-subdomains rotate relative to each other and to the C-subdomains (Flynn et al. 2003; Lewis et al. 1996; Mowbray and Björkman 1999).

**Figure 3.**
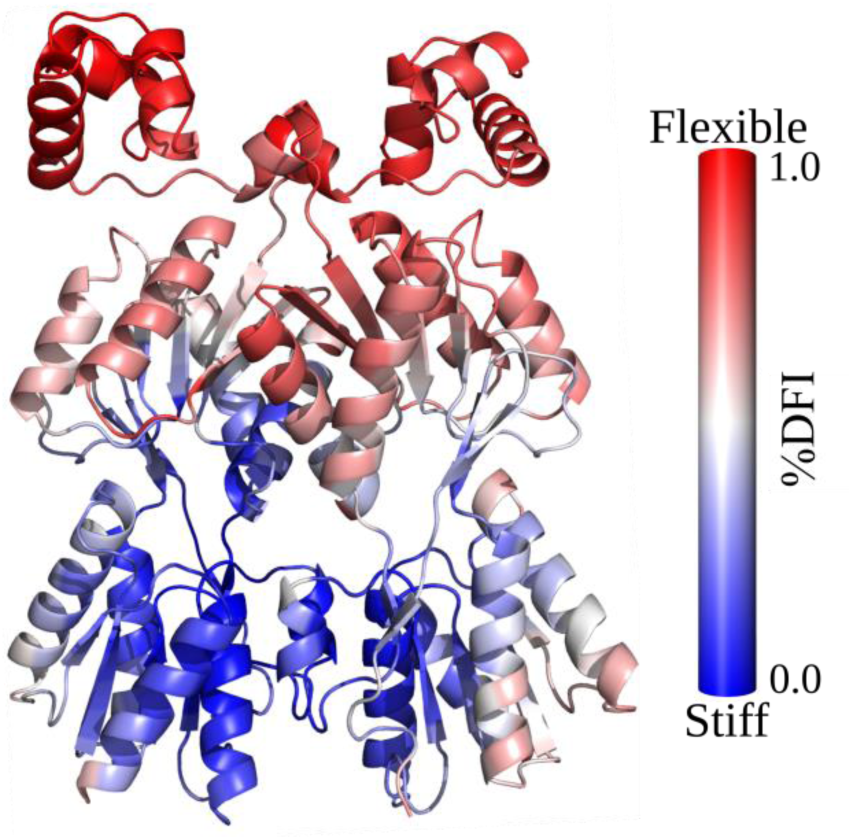
WT LacI (pdb ID 1efa (Bell and Lewis 2000)) structure colored by %DFI. The profile was calculated using each position on both chains from simulations of the DNA-bound structure in the absence of DNA or anti-inducer ONPF. Note that the DNA binding domains and the linker regions are more flexible than the distal parts of the regulatory domains, in agreement with known conformational changes.

### LacI V52X Variants Induce Local and Non-local Flexibility Changes in the DNA Binding Domain and Linker Region

Next, we determined whether amino acid substitutions in the LacI linker changed the flexibility profile of the DNA binding domain and whether the altered flexibility correlated with altered DNA binding affinities (K_d_ values). To that end, we first modeled the amino acid changes for eleven variants at position 52 and performed 250 ns molecular dynamics (MD) simulations with the AMBER ff14SB force field (D.A. Case et al. 2017). Relative to wild-type, the overall fold of the modeled structures remained unchanged for the DNA binding and regulatory domains (supplementary fig. S2A). However, the substitutions gave rise to differences in local interactions, as expected for a position that makes linker-linker interactions (supplementary fig. S2B and S2C) (Swint-Kruse et al. 2002), and subsequently we expected that these changes in local interactions would lead to changes in flexibility amongst the variants. Thus, to evaluate any flexibility changes, we calculated *%*DFI for positions in each of the V52X variants by using co-variance matrices obtained from MD simulations (see Methods).

Figure 4A shows the DNA binding domain and linker regions of wild-type LacI and three V52X variants colored by *%*DFI. Figures for the other variants are in supplementary fig. S3 and plots of %DFI values per position are shown in supplementary figs. S4 and S5. V52I had a similar *%*DFI profile in its DNA binding domain and linker region as wild-type LacI (fig. 4A), whereas seven variants with diminished DNA binding affinities exhibited stiffening throughout the linker region as well as varied and distinct changes in the DNA binding domain (V52T in fig. 4A and other variants in supplementary fig. S3). Finally, V52A – which exhibited the highest repression and enhanced DNA binding of the V52X variants in this study (table 1, supplementary fig. S1) – showed much more stiffening in both the DNA binding domain and linker.

**Figure 4.**
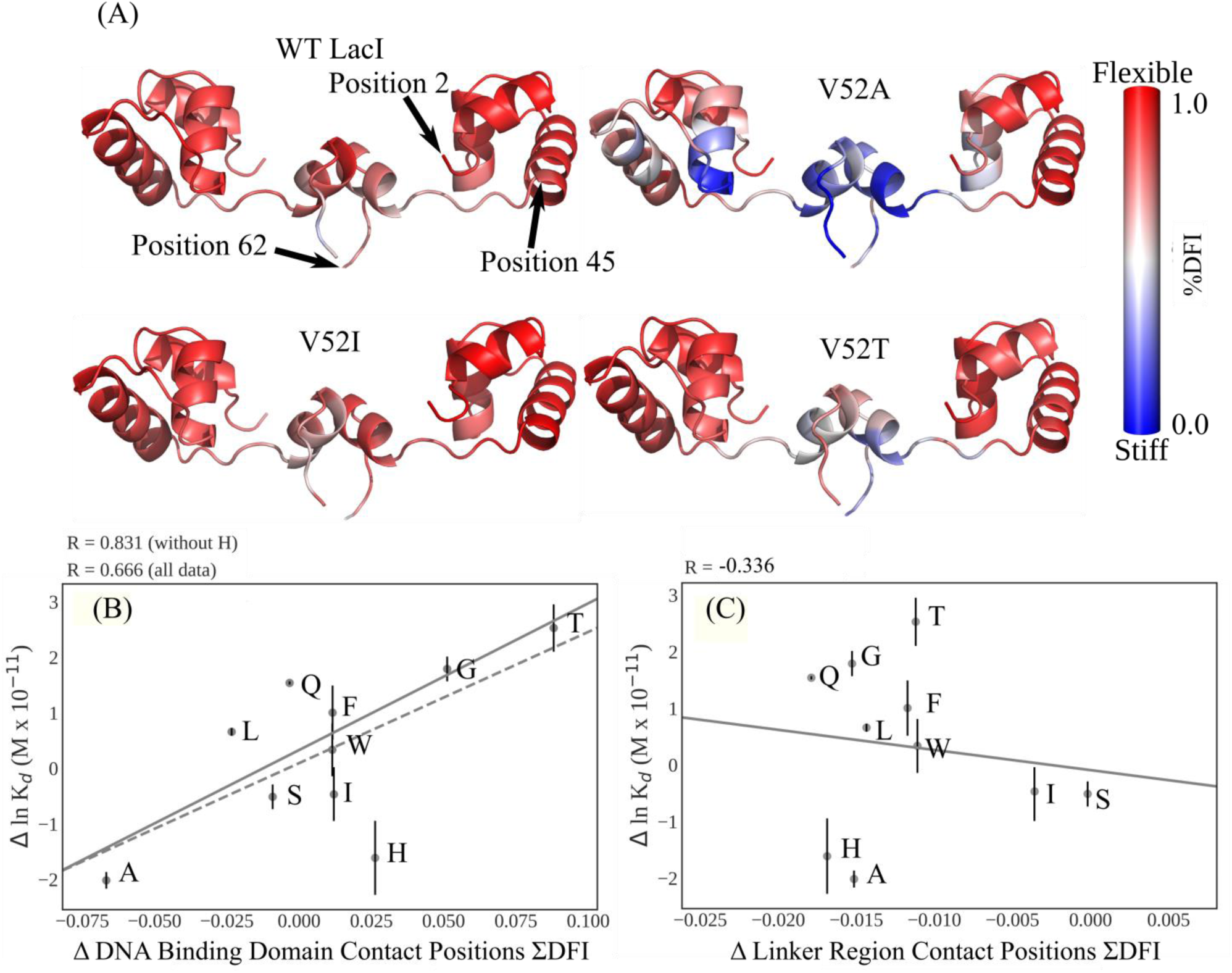
(A) V52X variants showed altered flexibility. The DNA binding domains (positions 2-45) and linker regions (positions 46-62) of LacI chains A and B are colored by %DFI for WT and three variants. All results are shown on the WT LacI crystal structure (pdb ID 1efa (Bell and Lewis 2000)). Note that V52A and V52T alter the flexibility profiles of DNA binding domain and linker region, whereas V52I preserves the WT flexibility. For variants not shown here, see supplementary fig. S3. Plots of %DFI per position are shown in supplementary figs. S4 and S5. (B, C) Change in DNA binding affinity vs change in ΣDFI for LacI positions that directly contact DNA. In the DNA binding domain, the direct contact positions include 3-7, 14-19, 21, 22, 24, 25, 28-32, and 34. In the linker region, the direct contact positions include 50, 53, 54, 56, 57, and 59. Calculations included positions on both monomers of a dimer. Black lines indicate the best fit from linear regression of the data. The DNA binding positions (B) show a strong correlation whereas no meaningful correlation exists between linker positions that contact DNA (C). Thus, we propose that V52X mutations lead to the long-distance changes in flexibility of DNA binding domain that modulate binding affinity. The V52H variant is a notable outlier on panel (A). One possibility arises from the fact that calculations were performed for a structure that is very similar to the DNA-bound structure used to initiate MD simulations. In contrast, changes in ln Kd can arise from changes in either the DNA-bound or unbound states. The V52X substitutions may also induce flexibility changes in the apo structure. This could be a contributing factor to both the V52H outlier as well as the imperfect correlation in (A).

Next, we reasoned that DNA binding affinities would emerge as a property of structural flexibilities in the DNA binding domain and possibly the linker region. The latter also contains residues that directly contact DNA (Bell and Lewis 2000; Kalodimos et al. 2001; Kalodimos et al. 2004; Kalodimos, Bonvin et al. 2002; Swint-Kruse et al. 2002). Therefore, we (i) summed the raw DFI values (“ΣDFI”) for various groupings of these positions and (ii) calculated the change in ΣDFI relative to that of wild-type LacI (“ΔΣDFI”). Finally, we correlated the changes in ΣDFI for various groups of positions with changes in measured DNA binding affinities (fig. 4B,C, supplementary fig. S6). Since DFI is a measure of local conformational entropy, it is appropriate to correlate this parameter with ln K_d_ values, which is proportional to free energy. Strikingly, for the DNA binding domain positions that directly contact DNA, enhanced flexibility correlated with diminished DNA binding affinity (fig. 4B). In contrast, no meaningful correlation was observed for linker positions that directly contact DNA (fig. 4C). Similar results were observed when all positions in the DNA binding domain and linker region were included in ΣDFI (supplementary figs. S6A, S6B, S6C). Thus, we conclude that scores for DNA binding domain positions with direct DNA contacts appeared contribute the most signal to the correlation.

### V52X Substitutions Alter Dynamic Coupling Between the Linker and DNA Binding Domains

The DFI results suggest that V52X substitutions have long-range repercussions on the flexibilities of the DNA binding domains. Thus, we used the dynamic coupling index (DCI) to quantify long-range dynamic coupling between position 52 and individual positions in the DNA binding domain (*e*.*g*. fig. 5A). The DCI parameter captures the strength of a displacement response for position *i* upon perturbation of position *j*, relative to the average fluctuation response of position *i* to all other positions in the protein. As such, this parameter represents a measure of the dynamic coupling between *i* and *j*. As in the case of %DFI, the DCI results are presented as %DCI, a percentile rank of the DCI range observed with values ranging from 0 to 1. Note that these two metrics are distinct: DFI is a measure of flexibility and DCI is a measure of coupling. Furthermore, DCI specifically quantifies the coupling between individual positions and, as such, DCI values depend explicitly upon the positions selected for analysis.

**Figure 5.**
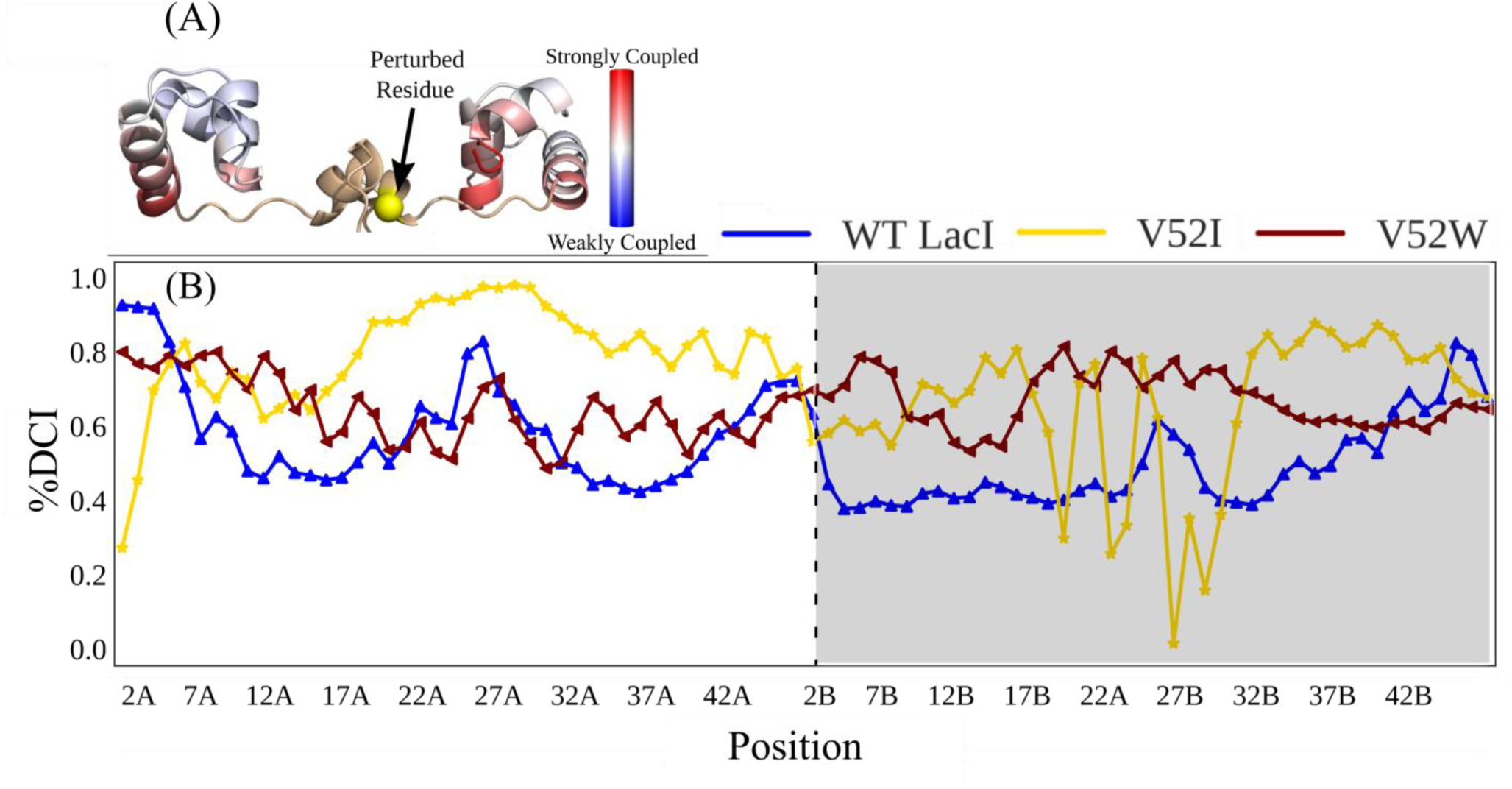
(A) Example DCI calculations mapped on the LacI DNA binding domain/linker structure. The DCI values for all DNA binding and linker positions that result from perturbing position 52 (yellow sphere) are shown with the red to blue color gradient. (B) Percentile-ranked dynamic coupling between position 52 of chain A and the DNA binding domain positions of WT LacI, V52W and V52I. The “A” and “B” designations on the position numbers and the white/gray shading indicate the two monomers of a dimer. Connecting lines are to aid visual inspection of the data. Note the significant differences in the score profiles, even though both variants have K_d_ values that are similar to WT. See supplementary fig. S7 for %DCI scores for other V52X variants.

For each V52X variant, we calculated %DCI values between position 52 of chain A and all positions in the two DNA binding domains (fig. 5B, supplementary fig. S7). Results show that each amino acid substitution led to dramatic dynamic coupling changes with many positions in the DNA binding domains. Further, the changes were amino acid specific. For example, even though wild-type, V52I, and V52W exhibited similar binding affinities (table 1) and similar flexibilities (fig. 4A, supplementary fig. S3), the three variants have significantly different coupling patterns with positions in the DNA binding domains (fig. 5B).

To assess whether any similarities exist among the coupling patterns of the V52X variants, we performed principle component analysis of the *%*DCI profiles. The singular value decomposition of each variant’s %DCI profile was first assessed by plotting the first two principle components (fig. 6A). Surprisingly, this analysis suggested a correlation with an additional aspect of LacI function: V52W, V52A, V52H, and V52S all cluster on the positive side of the first principle component; and DNA binding by these variants are *less* sensitive to allosteric regulation that occurs when inducer binds to the regulatory domains (fig. 6B). (Note that biochemically-measured values for V52I inducibility are not available and thus V52I is not represented in fig. 6B). To compare the top three principle components, we created a dendrogram (fig. 6C). In this analysis, observed clusters reflect groups with similar DNA binding K_d_ values (table 1). For example, four variants with K_d_ near or lower than wild-type LacI (V52A, V52S, V52H and V52I) are closely clustered (red oval).

**Figure 6.**
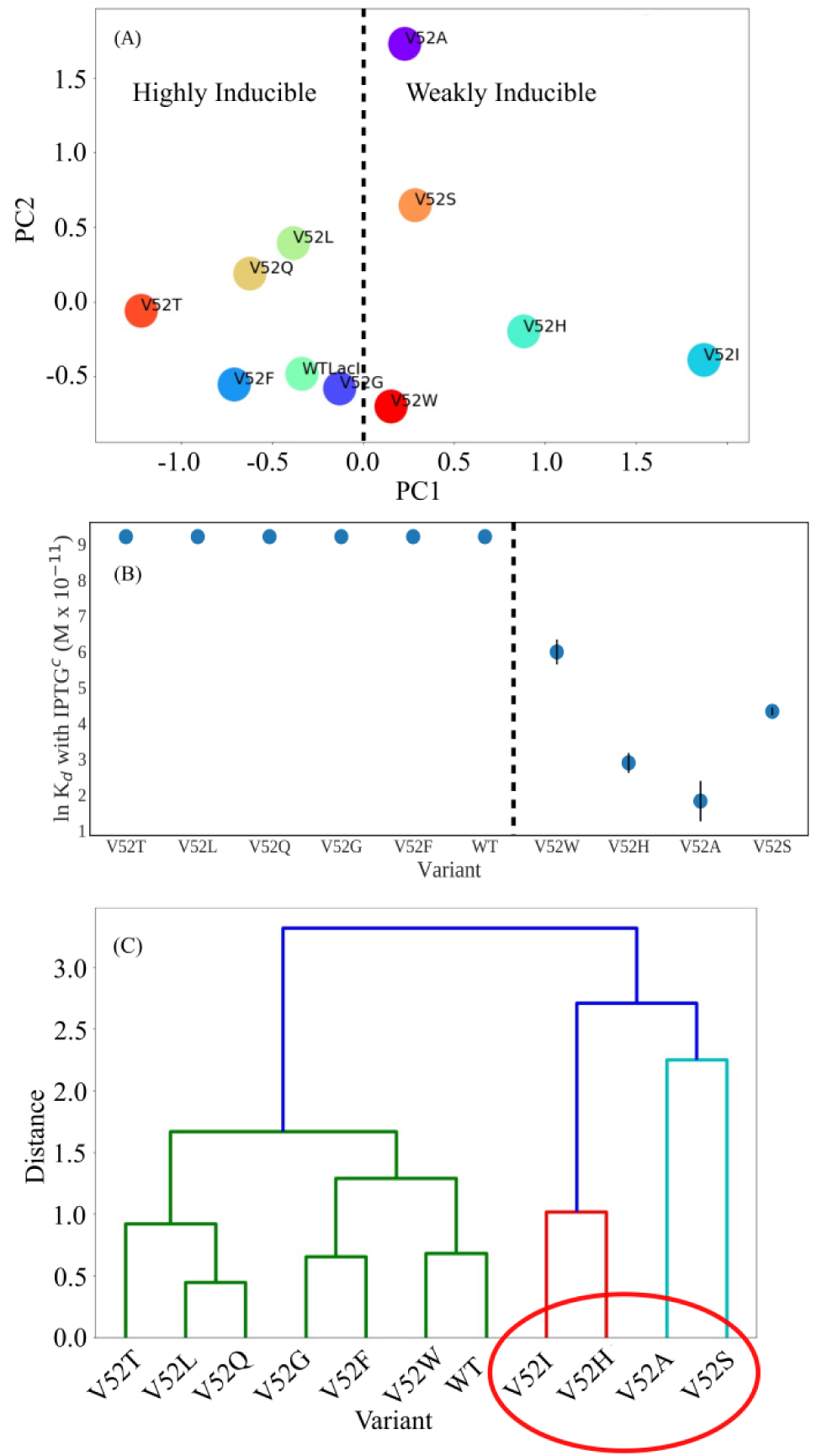
Principle component analyses of V52X %DCI scores. (A) Scatter plots of the first (PC1) vs second (PC2) principle components from the V52X to DNA binding domain %DCI scores suggest a separation in the first principle component correlates with altered inducibility (table 1), where the dashed line separates positive from negative values of the first principle component. Strikingly, this also appears to identify variants with altered inducibility (table 1 and panel (B)). (B) Point plot of ln K_d_ of DNA binding in the presence of IPTGc. The dashed line marks the separation of highly-inducible variants (K_d_ with IPTGc > 10000 M x 10-11) from weakly inducible variants and coincides with the dashed line in (A). The exception to this trend in PC1 may be V52I, which may comprise a third group as indicated by its position to the far right of panel (A). Biochemically-measured values for V52I inducibility are not available, but this variant showed WT inducibility in repression assays (Meinhardt et al. 2013). (C) Hierarchical clustering dendrogram using Ward’s method (Ward 1963) over first three principle components. Clusters associate with DNA binding K_d_ values both in the presence of IPTGc and without inducer (table 1). The colors represent clusters of variants within a given Ward distance cutoff used to define subgroups.

### Dynamic Coupling Asymmetries Can be Used to Distinguish Among Rheostat, Toggle, and Neutral Positions

Since the different V52X variants altered binding affinities across a wide range of values, we previously defined position 52 as a “rheostat” position (supplementary fig. S8, left panel) (Meinhardt et al. 2013). Rheostatic substitution behavior contrasts with the behavior often observed for conserved positions: For conserved positions in the LacI linkers, most substitutions are catastrophically deleterious (Markiewicz et al. 1994; Suckow et al. 1996) and we have referred to such positions as “toggle” positions (supplementary fig. S8, middle panel) (Hodges et al. 2018; Meinhardt et al. 2013; Miller et al. 2017). The LacI linker contains both toggle and rheostat positions (*e*.*g*. supplementary fig. S8) (Markiewicz et al. 1994; Meinhardt et al. 2013; Suckow et al. 1996; Zhan et al. 2006). Furthermore, the rheostat positions exhibited varied degrees of rheostat character, with some skewed more toward toggle behavior and others skewed more to “neutral” behavior (Hodges et al. 2018). At neutral positions, most substitutions have little to no effect; for example, LacI nonconserved position 62 is nearly perfectly neutral (supplementary fig. S8, right panel).

Since the DFI and DCI calculations predicted the functional outcomes for the individual substitutions at rheostat position 52, we were curious whether the wild-type structure alone could be used to predict the overall substitution outcome of each position. To that end, we used wild-type LacI to investigate (i) whether other positions with strong rheostatic characteristics showed similar coupling characteristics as position 52, and (ii) whether their characteristics of strongly rheostatic positions were distinct from positions that were dominated by toggle and neutral substitution outcomes. First, we performed two separate %DCI calculations: DNA binding domain responses to linker position perturbations and linker position responses to DNA binding domain perturbations. Neither of these methods distinguished the linker positions’ degree of rheostatic, toggle or neutral substitution behavior. However, we and others previously identified an intriguing characteristic in other model proteins (Campitelli et al. 2018; Guo et al. 2014; Hacisuleyman and Erman 2017; Schreiber 2000). In these systems, ligand binding altered the fluctuations at their local binding sites and modulated dynamics at distal sites via unidirectional communication. We reasoned that unidirectional communication might also be altered by amino acid substitutions and that analyzing the asymmetry in dynamic coupling could discriminate rheostatic or toggle substitution behavior.

As described in the Introduction, asymmetry can be captured in the DCI values because dynamic coupling is inherently asymmetric: For a particular pair of amino acids (*i* and *j*), their unique interactions constrain and differentiate the coupling of *i* to *j* from the coupling of *j* to *i* (*e*.*g*. fig. 2). Thus, we used the wild-type structure of LacI to calculate (i) %DCI_*i*_, how strongly each linker position is coupled to each DNA binding domain position, (ii) %DCI_*j*_, how strongly each DNA binding domain position is coupled to each linker position. From these, we calculated (iii) “%DCI_asym_” from (%DCI_*i*_ – %DCI_*j*_) to assess the asymmetry in coupling. Figure 7A shows %DCI_asym_ values obtained for positions representing the most rheostatic (V52), toggle-like (A53) and neutral (L62) positions; score distributions are presented as histograms in figures 7B-D. Results showed intriguing patterns, particularly when considering their coupling to positions 2-14 in the DNA binding domain. Values for the toggle position 53 were mostly negative, showing high unidirectionality, (the DNA binding domain was more strongly coupled to the linker region). On the other hand, values for the rheostat position 52 were less negative, whereas the values for neutral position 62 were near zero or slightly positive.

**Figure 7.**
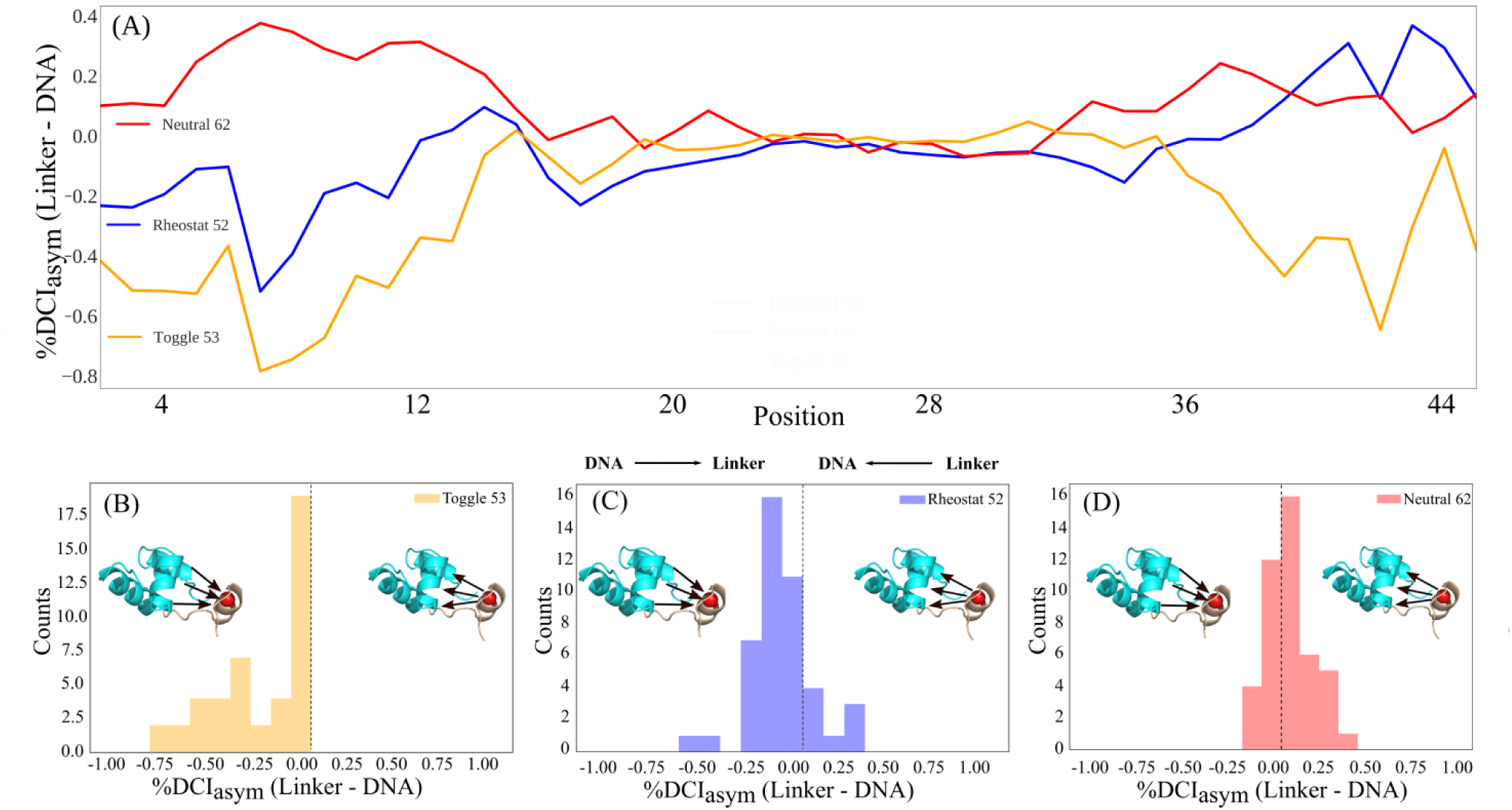
(A) %DCI_asym_ (linker to DNA binding domain - DNA binding domain to linker) scores for representative rheostat (52), neutral (62), and toggle (53) positions of chain A in WT LacI. (B-D) Distributions of %DCI_asym_ values for each position.

To assess whether these trends were broadly observed, we performed a principle component analysis of the %DCI_asym_ values for all of the linker rheostat, toggle, and neutral positions. Figure 8A shows a plot of the first two principle components from this decomposition. The indicated groups were empirically determined in the *absence* of detailed knowledge about functional outcomes for each position. When functional data (fig. 8B and supplementary fig. S9) were compared to computational results,, PC2 appeared to separate the toggles from non-toggle positions (supplementary fig. S10). When additional analyses were conducted for non-toggle positions using PC1, PC3, PC4 and PC5, the resulting dendrogram (supplementary fig. S11A) as well as a direct comparison of PC1 and PC3 (supplementary fig. S11B) suggest that the non-toggle positions form clusters that are similar to groups shown in figure 8A.

**Figure 8.**
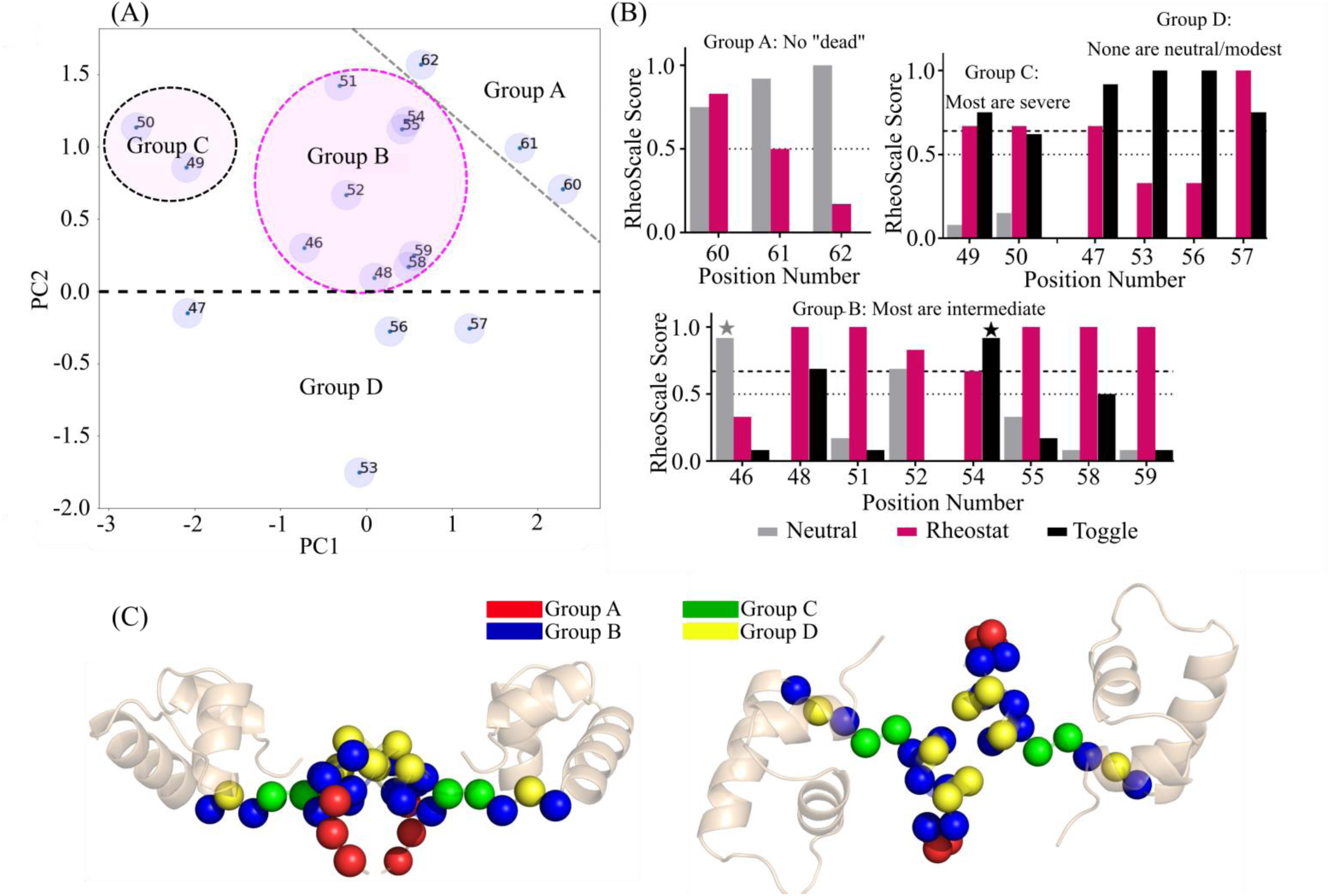
Comparison of %DCI_asym_ analyses with functional attributes of each liker position. (A) 2-D plot of the first two principle components from singular value decomposition of %DCI_asym_ profiles. Four groups were identified and indicated on this plot. (B) Aggregate of the experimental outcomes for all substitutions at each LacI linker position. For each position, the outcomes from all available substitutions were analyzed with the “RheoScale” calculator (Hodges et al. 2018) to describe each position’s overall substitution behavior. “Neutral” scores (gray bars) reflect the fraction of substitutions that have little effect on function. “Rheostat” scores (magenta bars) reflect the fraction of the functional range that was accessed by at least one substitution; for the high-resolution dataset, we previously defined rheostat scores ≥0.5 (dotted line) as significant (Hodges et al. 2018). “Toggle” scores (black bars) reflect the fraction of substitutions that are greatly damaging to function (“dead”); we previously defined toggle scores ≥0.67 (dashed line) as significant (Miller et al. 2017). This panel shows calculations for the low-resolution experimental data, which are available for all linker positions; calculations for the high-resolution data, which are available for 11 linker positions, are shown in supplementary fig. S9; for further details about these two datasets, see Methods. Note that the neutral and toggle scores determined from low resolution assay data are over-estimates; many of these substitutions have intermediate outcomes. Scores for two positions affected by this over-estimate are indicated with gray and black stars. (C) Linker position Cα carbons colored by cluster for face-forward (left) and top-down (right) views

When the experimental substitution behaviors were compared to the principle component analyses (fig. 8B and supplementary fig. S9), the groups from the %DCI_asym_ analysis matched reasonably well with the groups derived from experimental data (Hodges et al. 2018): For the three positions in group A, no catastrophic (“dead”) substitutions were identified in either of two experimental studies; instead most substitutions had neutral or modest outcomes. Positions with strong rheostat scores – which indicates that many intermediate outcomes were observed – fell into group B (see also supplementary fig. S9). For the two positions in group C, most substitutions were catastrophic, but a few substitutions showed modest or neutral effects. Finally, for positions in group D, neither study identified modest or neutral substitutions and the majority of substitutions were catastrophic (as indicated by high toggle scores and neutral scores of zero). Figure 8C shows the individual positions of each group from 8A and 8B color-coded on the LacI structure. Interestingly, the positions in groups A and D generally exhibited %DCI_asym_ distributions that were similar to those shown for example toggle and neutral positions in figure 7D and 7B.

In summary, %DCI_asym_ analysis was able to largely discriminate the different types of substitution outcomes experimentally observed for the LacI linker positions. The toggle-like and the neutral-like positions each comprised one group, and the rheostat positions were divided into two groups with different maximal severities from substitution.

## Discussion

For missense variants, the prediction accuracy of current diagnostic tools is astonishingly low. In addition to their participation in multi-factorial diseases, one source of the low accuracy arises from variants at nonconserved positions (the fastest evolving positions) (Miller et al. 2017; Subramanian and Kumar 2006). These positions are often *not* in close proximity to functionally critical sites (Curado-Carballada et al. 2019; Jiménez-Osés et al. 2014; Modi and Ozkan 2018; Saavedra et al. 2018; Townsend et al. 2015), which has made it difficult to rationalize how their substitutions alter function. Similar observations have been made in directed evolution studies, which frequently identify substitutions distal from active, conserved sites that are required to evolve functional variation (Swint-Kruse 2016; Wilding et al. 2019).

In attempts to understand the effects of missense variants in the LacI linker, extensive correlations with analyses of multiple sequence alignments were previously employed (Meinhardt et al. 2013; Parente et al. 2015; Parente and Swint-Kruse 2013; Tungtur et al. 2011). The nonconserved LacI linker positions do not have strong co-evolution scores, but they have reasonably high phylogeny scores. Since the current data is limited to the linker region of LacI, the number of positions is too small to determine a definitive correlation. Work is ongoing to determine whether there is a general relationship between the toggle, neutral, and rheostat positions and scores from various sequence analyses (including sequence entropy). In addition, both phylogeny and/or co-evolution scores have been assessed for other types of structural or functional correlations in various LacI/GalR homologs, such as ligand specificity or direct structural contacts; to date, no relationship has been found (Meinhardt and Swint-Kruse 2008; Parente and Swint-Kruse 2013; Taraban et al. 2008; Tungtur et al. 2010; Tungtur et al. 2019).

Here, we have analyzed whether protein structural dynamics can provide insights how substitutions at nonconserved sites modulate protein function. Using LacI as a model system, with specific focus on the relationship between the linker region and DNA binding domain, our calculations showed correlations with experiment for individual substitutions at one nonconserved linker position and could be used to discriminate the overall substitution responses for linker positions with a range of substitution profiles. Results provide mechanistic understanding to LacI function, suggest avenues for future computations to further understand the substitution/function relationship in LacI, and suggest trends that may be generally applicable to other proteins.

### DFI and DCI Calculations Inform Several Aspects LacI Function

Experimentally, when apo-LacI binds to DNA, the protein undergoes a large conformational change (Ha et al. 1989; Kalodimos, Boelens et al. 2002; Swint-Kruse et al. 2002; Taraban et al. 2008). The DFI calculations (fig. 4 and supplementary fig. S3) suggest that V52X substitutions, and perhaps substitutions for other linker rheostat positions, alter the equilibrium dynamics of the DNA bound conformation. In turn, these entropic changes would alter the free energy of DNA binding that was captured in experimental measures of DNA binding affinity. Although not considered in the current computations, the V52X substitutions might also alter the equilibrium dynamics of the unbound state or induced states. Differential changes in each of the conformations may help explain why the correlations between binding affinity and DFI were not perfect (fig. 4B). Changes in flexibility were calculated on DNA-bound ensembles only, whereas the experimentally-determined free energy of binding is defined by the difference between bound and unbound states. In future studies, the predicted entropic changes could be tested using thermodynamic experiments designed to detect the entropic component of DNA binding.

In addition to DNA binding, LacI binds inducer molecules via its regulatory domain (fig. 1), and the DNA binding and regulatory domains exhibit allosteric communication (reviewed in (Swint-Kruse and Matthews 2009)). The latter function was also altered by substitutions at position 52 (Zhan et al. 2006). Interestingly, clustering the dynamic coupling index (DCI) of the V52X positions captured both the changes in the DNA binding and the changed response to inducer (Zhan et al. 2006) (fig. 6). Analogous to the DNA binding positions that showed strong correlation in figure 4B, we speculate that a group of “sensor” positions in the inducer binding site will show long-range flexibility changes in response to amino acid changes at position 52. In addition to the DNA bound conformation used herein, such computations should be pursued using the inducer-bound conformation of LacI. It will also be interesting to determine whether DCI/DFI calculations can be used to predict the locations of other nonconserved positions for which substitutions modulate LacI-inducer binding or allosteric regulation. More generally, this observation suggests a mechanism by which substitutions change in long-distance dynamic coupling to modulate allosterically regulated proteins.

In addition to alternative protein conformations, alternative DNA sequences might contribute to computational outcomes. For this work, the LacI structure was extracted from a structure that was bound to DNA. During the simulations, LacI did not transform all the way into the apo-form: the hinge helices in the linker stayed folded and the DNA binding domains maintained similar conformations (supplementary fig. S1C). Thus, some of the information from the DNA sequence may have persisted during the simulations and DFI/DCI calculations. In the crystal structure used for this work (pdb 1efa), LacI was bound to the symmetric operator “Osym” (Bell and Lewis 2000). However, the experiments referenced herein used the natural operator “O1” (Meinhardt et al. 2013; Zhan et al. 2006). This did not concern us, because the effects of most V52X variants are comparable for these two DNAs (Tungtur et al. 2019). (When a substitution enhanced Osym binding, O1 binding was enhanced by a comparable amount). However, this relationship between Osym and O1 binding is not known for V52I, which is an outlier in figure 6A. Thus, one prediction could be that V52I-Osym binding in the presence of inducer may differ from the trend followed by the rest of the V52X variants.

### DCI_asym_ Distributions Capture Substitution Outcomes Trends at Rheostat, Toggle, and Neutral Positions

When we expanded our calculations to consider the dynamic coupling between 17 linker positions with the DNA binding domain in the wild-type protein, we observed that degree of asymmetry in long range dynamic coupling was in agreement with the functional outcomes of substitution. As discussed above, the dynamic coupling between linker position *i* and the DNA binding position *j* is not necessarily symmetric, due to the asymmetry inherent to each positions’ local network of interactions. This asymmetry gives rise to unidirectional information transfer, a concept which has previously been used to identify asymmetry in allosteric systems and to predict mutational impact (Hacisuleyman and Erman 2017; Kamberaj and van der Vaart 2009; LeVine and Weinstein 2014).

Historically, we have considered the “obvious” explanations for the origins of toggle positions. For example, LacI toggle position A53 directly contacts DNA; most other substituted side chains might make a steric clash; the A53G substitution (also catastrophic) might lack a critical interaction. The current work provides another explanation – not only does position 53 participate in the DNA interaction, it is a key node in the LacI coupling network. Indeed, all four toggle-like positions in the linker may function as key nodes in the coupling network. These positions exhibited large coupling asymmetry, with greater dynamic coupling of the DNA binding domain to the linker region than the reverse. In other words, all the DNA binding domain positions dominated the long-range communications to these linker positions, suggesting that substitution at these positions could compromise these critical communication channels with the DNA binding domain. In contrast, the three neutral-like positions in the linker exhibited largely symmetric distributions, with a balance between DNA-dominant and linker-dominant coupling.

Intriguingly, the distributions of %DCI_asym_ values for rheostat linker positions were intermediate to the two extremes (toggle and neutral). This may allow (or be requisite for) broad modulation of function: Substitutions at rheostat positions may need only to slightly alter the dynamic coupling/communication to increase or decrease dynamic coupling among distal positions, (*i*.*e*. re-wire a new dynamic coupling network). This would explain the long-range effects of substitutions and gives rise to another intriguing possibility for the underlying basis of non-additivity often observed among two or more substitutions. Finally, this dynamic coupling network provides a satisfactory framework for understanding the biophysical mechanisms that underlie protein evolution and could explain why many of the rheostat positions identified to date occur at nonconserved positions.

### Conclusion

In summary, these calculations suggest a number of promising avenues for understanding the functional outcomes of amino acid substitutions, particularly at nonconserved sites. These results add to the growing body of evidence that substitutions at nonconserved sites modulate function by fine-tuning flexibilities of distal functional sites. Modulation of flexibilities was also observed in studies that compared the equilibrium dynamics of resurrected ancestral proteins to modern-day homologs (Kim et al. 2015; Modi and Ozkan 2018; Zou et al. 2015). Similarly, long-range dynamics regulation and altered DCI profiles were observed in evolutionary variants of thioredoxin (Modi et al. 2018) and disease-associated human missense variants (Kumar et al. 2015). Thus, one mechanism by which proteins can adapt a new environment or acquire a new function (*i*.*e*. degrade a new substrate) is *via* substitutions that change the flexibilities of functionally-important distal sites (Butler et al. 2015; Curado-Carballada et al. 2019; Hilser 2010; Jiménez-Osés et al. 2014; Modi and Ozkan 2018; Saavedra et al. 2018; Townsend et al. 2015). If the change is too large, the extreme change manifests as disease.

Studying the changes in equilibrium dynamics and, particularly, analyzing how substitutions modulate conformational states could provide mechanistic insights as to how substitutions at nonconserved sites affect function. In the future, this type of analysis can be expanded to other regions of LacI and to other types of proteins. In combination with dynamic spectroscopic analysis (single molecule or ensemble), computational mapping of the full interaction networks, particularly those existing between functional domains, will allow clear identification of critical positions in these networks, such as multi-pathway nodes. Such analyses could provide a means to predict the severity of substitution outcomes at non-conserved positions

## Methods

### Functional data for the LacI V52X variants

As described in figure 1, a dimer is the minimal LacI functional unit for high affinity DNA binding. However, full-length wild-type LacI is a dimer-of-dimers, with tetramerization facilitated by an additional C terminal domain that forms a coiled coil. This domain can be cut off, leaving dimer with essentially identical intrinsic binding affinity for a single DNA binding site (Chen and Matthews 1994). DNA binding affinities reported in table 1 were measured for the tetrameric versions of LacI (Zhan et al. 2006). High resolution repression values (table 1) (Meinhardt et al. 2013) were measured using a dimeric version of LacI in order to compare high resolution *in vivo* repression to that other dimeric paralogs (repression by tetrameric LacI is enhanced by binding and two looping two operator sequences) (Swint-Kruse and Matthews 2009). In all studies carried out to date, substitution outcomes on DNA binding affinities are the same in tetrameric and dimeric LacI (Swint-Kruse et al. 2005; Tungtur et al. 2019). This is supported by the strong correlation shown in fig. S2. In this manuscript, we use the term “wild-type” interchangeably for the dimeric and tetrameric forms. A third dataset comprising low-resolution *in vivo* repression data for variants of tetrameric LacI (Markiewicz et al. 1994; Suckow et al. 1996) was used for RheoScale analyses and is further described below.

In both biochemical and high-resolution *in vivo* studies, all substitution outcomes appeared to manifest in functional changes, rather than as changes in structure or stability: In the *in vitro* study, the variant proteins purified and behaved very similar to wild-type LacI (Zhan et al. 2006). In the *in vivo* studies, control experiments showed that all protein variants were expressed and capable of binding DNA (Meinhardt et al. 2013). Together, these observations indicate that the gross structural features of the LacI V52X variants were not distorted.

### Modeling of V52X variant structures

All LacI V52X variants were modeled using chains A and B from the structure of the dimeric form of the *E. coli* Lac repressor (Bell and Lewis 2000) (PDB ID 1efa); the models for DNA operator and anti-inducer ONPF were removed prior to calculations. Mutagenesis was performed using the PyMol Mutagenesis Wizard on valine at position 52 of both chains A and B to create 10 additional variants: V52A, V52F, V52G, V52H, V52I, V52L, V52Q, V52S, V52T and V52W. Topology files for all structures were prepared using the AMBER LEaP program with the ff14SB force field (D.A. Case et al. 2017), where hydrogen atoms were added and each structure was surrounded by an 18.0 Å cubic box of water molecules using the TIP3P (MacKerell et al. 1998) water model. Na^+^ and Cl^-^ atoms were added for neutralization. To remove any unfavorable torsional angles or steric clashes and ensure that the system reached a local energetic minimum, each system was energy-minimized using the AMBER SANDER package (D.A. Case et al. 2017). First, the protein was kept fixed with harmonic restraints to allow surrounding water molecules and ions to relax, followed by a second minimization step in which the restraints were removed and the protein-solution was further minimized. Both minimization steps employed the method of steepest descent followed by conjugate gradient.

The systems were then heated from 0K to 300K over 250 ps, at which point long-range electrostatic interactions were calculated using the particle mesh Ewald method (Darden et al. 1993). Direct-sum, non-bonded interactions were cut off at distances of 9.0 Å or greater. The systems were then simulated using Molecular Dynamics at constant temperature and pressure with 2fs time steps for 250 ns. Covariance matrix data were calculated over the final 40 ns of each simulation, using a 20 ns moving window that overlap by 5 ns. Additional references used to validate these models were (Swint-Kruse et al. 1998; Swint-Kruse et al. 2001; Swint-Kruse 2004; Swint-Kruse and Brown 2005), as further detailed in the supplement.

### Dynamic Flexibility and Dynamic Coupling

The Dynamic Flexibility Index utilizes a Perturbation Response Scanning technique that combines the Elastic Network Model and Linear Response Theory (“LRT”) (Gerek and Ozkan 2011; Nevin Gerek et al. 2013)

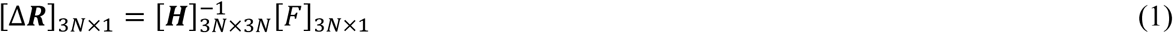

Using LRT, Δ***R*** is calculated as the fluctuation response vector of residue *j* as a result of unit force’s ***F*** perturbation on residue *i*, averaged over multiple unit force directions to simulate an isotropic perturbation (2). ***H*** is the Hessian, a 3N × 3N matrix which can be constructed from 3-D atomic coordinate information and is composed of the second derivatives of the harmonic potential with respect to the components of the position’s vectors of length *N*. For this work, the Hessian matrix was extracted directly from molecular dynamics simulations as the inverse of the covariance matrix. This method allows one to implicitly capture specific physiochemical properties and more accurate residue-residue interactions via atomistic force fields and subsequent all-atom simulation data.

Each position in the structure was perturbed sequentially to generate a Perturbation Response Matrix ***A***

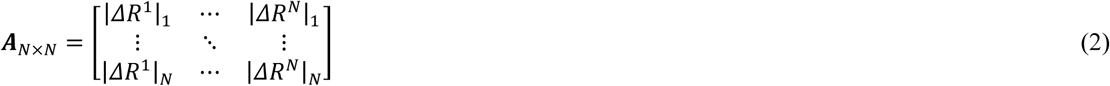

where 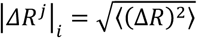 is the magnitude of fluctuation response at position *i* due to perturbations at position *j*. The DFI value of position *i* is then treated as the displacement response of position *i* relative to the net displacement response of the entire protein, which is calculated by sequentially perturbing each position in the structure (3).

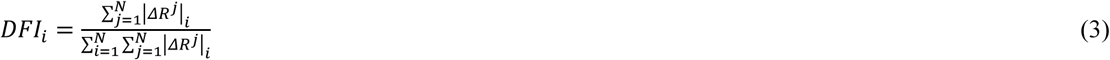

It is also often useful to quantify position flexibility relative to the flexibility ranges unique to individual structures. To that end, DFI can be presented as a percentile rank

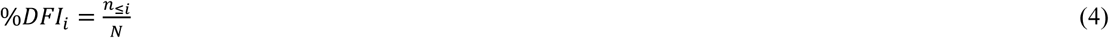

where *n*_*≤i*_ is the number of positions with a DFI value ≤ DFI_i_. The denominator is the total displacement of all residues, used as a normalizing factor. All %DFI calculations present in this work used the DFI value of every residue of the full LacI structure for ranking. The DFI parameter can be considered a measure of a given amino acid position’s ability to explore its local conformational space.

### Dynamic Coupling Index

Similar to DFI, the dynamic coupling index (DCI) also utilizes Perturbation Response Scanning with the Elastic Network Model and Linear Response Theory. The dynamic coupling index (DCI) captures the strength of displacement response of a given position *i* upon perturbation to a single functionally important position (or subset of positions) *j*, relative to the average fluctuation response of position *i* when all of the positions within a structure are perturbed.

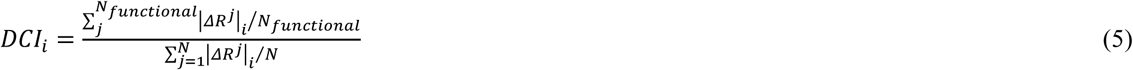

As above, DCI can also be presented as a percentile rank as

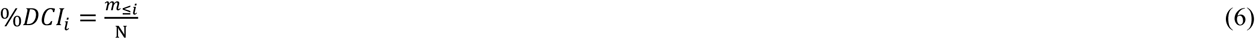

where *m*_*≤i*_ is the number of positions with a DCI value ≤ DCI_i_. As such, this parameter represents a measure of the dynamic coupling between *i* upon a perturbation to *j*. Additional information such as preferential information transfer through asymmetric dynamic coupling can also be determined by analyzing the coupling asymmetry between positions *i* and *j*, which can be calculated as

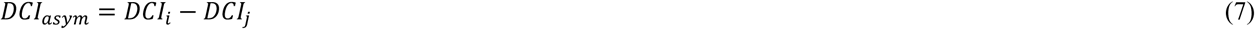

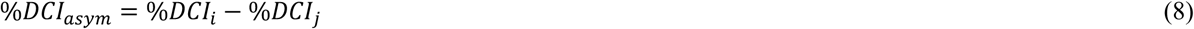

where a positive DCI_asym_ value indicates communication from position *i* to position *j*. For this work, DCI analysis was performed using force perturbations to chain A positions.

### RheoScale Calculations Aggregate Experimental Functional Data for Multiple Variants

Rather than thinking about the role of individual amino acid residues within a protein, we have found it useful to consider the evolutionary role of each protein position: Conserved positions often require a specific amino acid to carry out an evolutionarily conserved function (or structure). Most amino acid substitutions at these locations are catastrophic to the protein’s structure or function. As such, the outcomes for multiple substitutions resemble a toggle switch, with function either “on” or “off”. Alternatively, some nonconserved positions can be varied to modulate function, an evolutionarily advantageous method to help organisms adapt to new niches. We have shown that substitutions at these positions can lead to a wide range of intermediate outcomes (Hodges et al. 2018; Meinhardt et al. 2013). Finally, some positions can tolerate any amino acid substitution; functional outcomes are “neutral” (Martin et al. 2019; Miller et al. 2019).

These three categories represent idealized behaviors. Substitution data for real protein positions often fall on the continuum. Thus, we developed a scale to quantify how toggle-like, rheostatic, and neutral the aggregate functional data are for a given position (Hodges et al. 2018). “Neutral” scores reflect the fraction of substitutions that have little effect on function. “Rheostat” scores reflect the fraction of the functional range that was accessed by at least one substitution; for the high-resolution dataset, we previously defined rheostat scores ≥0.5 (dotted line) as significant (Hodges et al. 2018). “Toggle” scores reflect the fraction of substitutions that are greatly damaging to function; we previously defined toggle scores ≥0.67 (dashed line) as significant (Miller et al. 2017).

For the LacI linker positions, two datasets are available for RheoScale calculations, both comprised of *in vivo* repression measurements: (i) The Swint-Kruse lab used a high-resolution functional assay to assess outcomes for 7-14 substitutions at each of 11 nonconserved linker positions in dimeric LacI (Meinhardt et al. 2013). High resolution data for position 52 are used in table 1 and Rheoscale calculations for position 52 and other nonconserved positions (Hodges et al. 2018) are shown in supplementary fig. S9. (ii) The Miller lab used a low resolution assay to assess outcomes for 12-13 substitutions at all linker positions (and most other positions) in tetrameric LacI (Markiewicz et al. 1994; Suckow et al. 1996); calculations from these data are shown in supplementary fig. S8 and more details are given below. In all LacI studies to date, substitutions have the same outcome on DNA binding for tetrameric and dimeric LacI (*e*.*g*. (Swint-Kruse et al. 2005)). The two functional studies also have some differences in their sets of substituted amino acids used, but where overlapping data is available, the RheoScale scores were in good agreement (Hodges et al. 2018).

RheoScale calculations for the low resolution data were as follows: For each substitution, repression was scored as either (i) “+” (reporter gene activity, relative to “no repression”, was more than 200-fold diminished); (ii) “+/-” (diminished from 200 to 20 fold); (iii) “-/+” (diminished from 20 to 4 fold); or (iv) “-” (no more than 4-fold diminished) (Markiewicz et al. 1994; Suckow et al. 1996). Since the RheoScale calculator requires numerical data to perform histogram analyses, we reassigned these categories as numerical values, with 1 being the best repression and 4 being the worst. Histogram analyses were carried out using 4 bins. Note that the “+” data encompasses some substitutions with modest changes in repression that would be detected in the low-resolution assay; the “+” data also encompass any substitutions that enhance repression. Likewise, the “-” substitutions encompass some positions that retain some weak ability to repress. Thus, both the neutral and the toggle scores were over-estimated for these data, whereas the rheostat score was underestimated.

## Acknowledgements

This work was supported by the National Science Foundation Division of Molecular and Cellular Biosciences (award 1715591), the National Institute of Health (grant number GM118589), the Gordan and Betty Moore Foundation and the W.M. Keck Foundation.

