## supplementary for "Substitutions at Non-Conserved Rheostat Positions Modulate Function by Re-Wiring Long-Range, Dynamic Interactions"

**Table S1:** Shannon sequence entropies for the linker positions in alignments of LacI/GalR families

**Figure S1:** Experimentally determined changes in repression vs changes in DNA binding affinity for six V52X mutants (black letters).

**Figure S2:** Structural comparison and model validation of V52X variants.

**Figure S3**: DNA binding domain and linker region flexibility of V52X variants.

**Figure S4**: %DFI for DNA binding domain and linker region positions of chains A and B in WT and five V52X variants.

**Figure S5**: %DFI for DNA binding domain and linker region positions of chains A and B in remaining five V52X variants.

**Figure S6**. Changes in DNA binding versus changes in flexibility for various subsets of LacI positions.

**Figure S7**. %DCI scores between position 52 of chain A and all DNA binding domain positions in each of the V52X variants.

**Figure S8.** Example positions with rheostat, toggle, and neutral substitution outcomes.

**Figure S9**. RheoScale scores from high resolution functional data of each linker position. **Figure S10**. Linker positions plotted in order of increasing second principle component value from decomposition of DNA-Linker %$DCI_{asym}$.

**Figure S11.** Dendrogram (A) and PC3 vs PC1 point plot (B) of %$DCI_{asym}$ principle components excluding toggle positions (47, 53, 56, 57).

| Linker Position | Whole family | “YPAL” half-family |
|---|---|---|
| 46 | 1.56 | 1.57 |
| **47** | **0.24** | **0.27** |
| 48 | 2.25 | 2.12 |
| **49** | **0.7** | **0.15** |
| 50 | 1.2 | 0.93 |
| 51 | 2.15 | 2.06 |
| 52 | 2.24 | 1.93 |
| **53** | **0.91** | **0.12** |
| 54 | 1.37 | 0.89 |
| 55 | 2.21 | 1.91 |
| **56** | **0.96** | **0.53** |
| 57 | 1.98 | 1.73 |
| 58 | 2.37 | 2.23 |
| 59 | 2.14 | 1.8 |
| 60 | 2.28 | 2.21 |
| 61 | 1.68 | 1.35 |
| 62 | 2.3 | 2.2 |

**Supplemental Table 1. Shannon sequence entropies for the linker positions in alignments of LacI/GalR families.** Sequence entropies range from 0.00 (perfectly conserved) to 2.82 (perfectly random). Data were taken from (Tungtur et al. 2011). Four linker positions are highly conserved (bold): Linker position 47 is highly conserved in all LacI/GalR homologs, which comprises >34 subfamilies. Linker positions 49, 53, and 56 are highly conserved in the ~2/3 of LacI/GalR sequences (“half-family”) that have a “YxPxxxAxxL” motif at these same conserved positions (Tungtur et al. 2011). All other linker positions are nonconserved, to different degrees, in the LacI/GalR family.

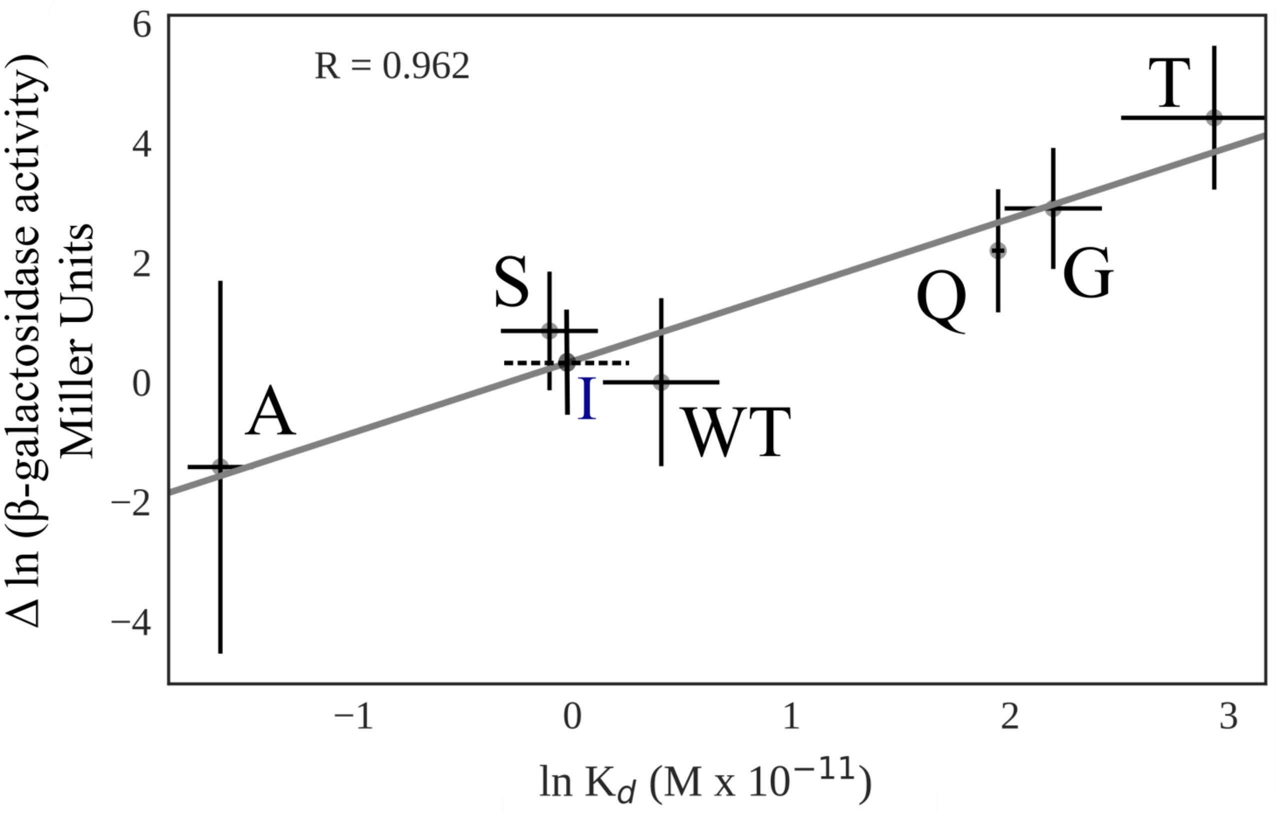


**Figure S1**. **Experimentally determined changes in repression vs changes in DNA binding affinity for six V52X mutants (black letters).** Here and in subsequent figures, the letter next to each data point indicates the amino acid change for position 52 (collectively referred to in this manuscript as "X"). "WT" indicates the data for wild-type LacI. Repression data are from (Meinhardt et al. 2013) and binding affinity data were taken from (Zhan et al. 2006); details of these two studies are further described in Methods. Error bars represent standard deviations of mean values. The solid line represents the best fit of the data by linear regression. The strong correlation observed between the two parameters indicates that changes in repression were largely due to changes in DNA binding affinities. The value and error for V52I DNA binding affinity (dashed error bar) was extrapolated from its repression value (Meinhardt et al. 2013) using linear regression.

(A)

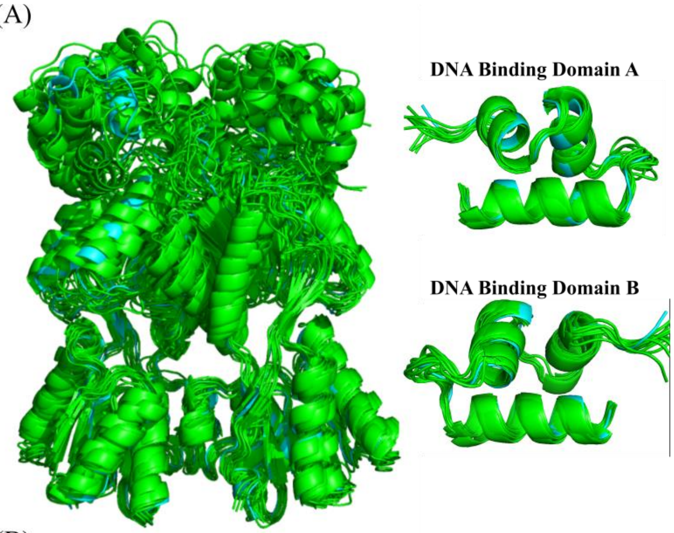


| RMSD (Å) | | | |
|---|---|---|---|
| Variant | Full Structure | DNA A | DNA B |
| WT | 2.48 | 0.61 | 0.87 |
| V52A | 1.55 | 0.85 | 0.84 |
| V52I | 1.87 | 0.58 | 0.55 |
| V52T | 1.44 | 0.77 | 1.06 |
| V52W | 1.5 | 0.86 | 0.71 |
| V52F | 1.76 | 0.96 | 0.76 |
| V52G | 1.49 | 0.63 | 1.12 |
| V52H | 2.16 | 0.86 | 1.22 |
| V52L | 2.09 | 0.76 | 0.61 |
| V52Q | 2.59 | 0.68 | 0.53 |
| V52S | 1.87 | 0.59 | 0.96 |

(B)

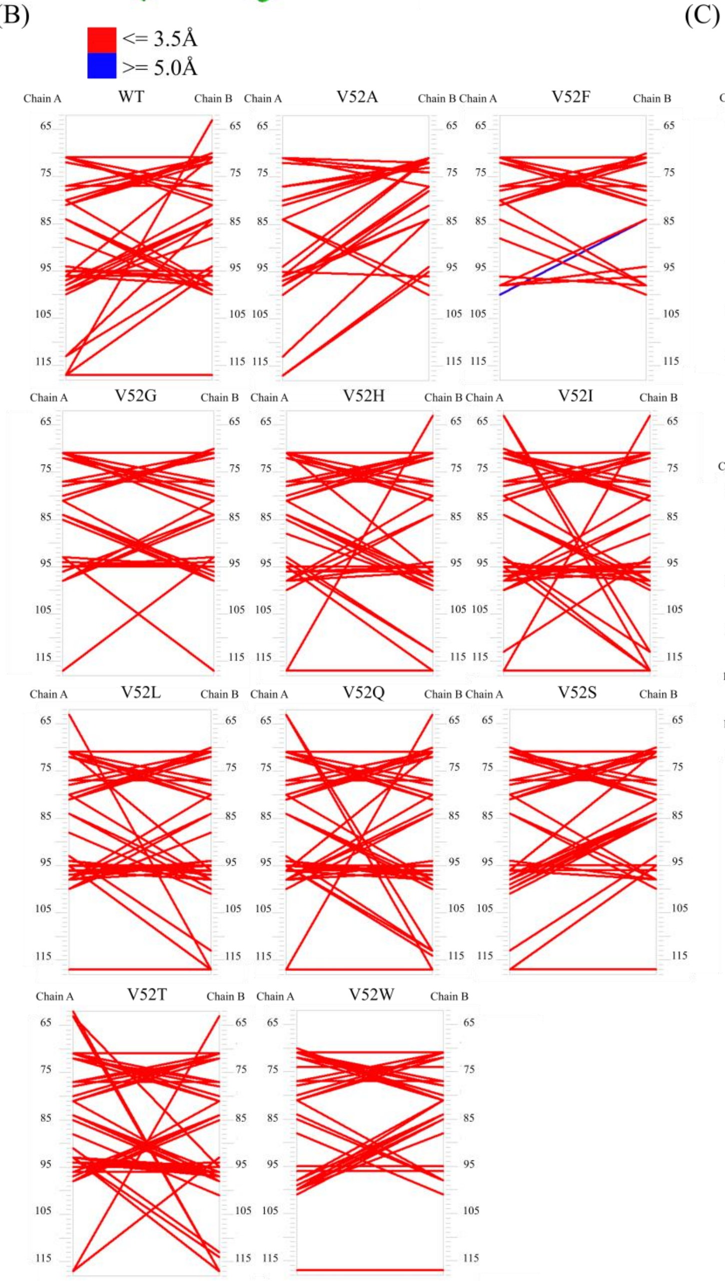


(C)

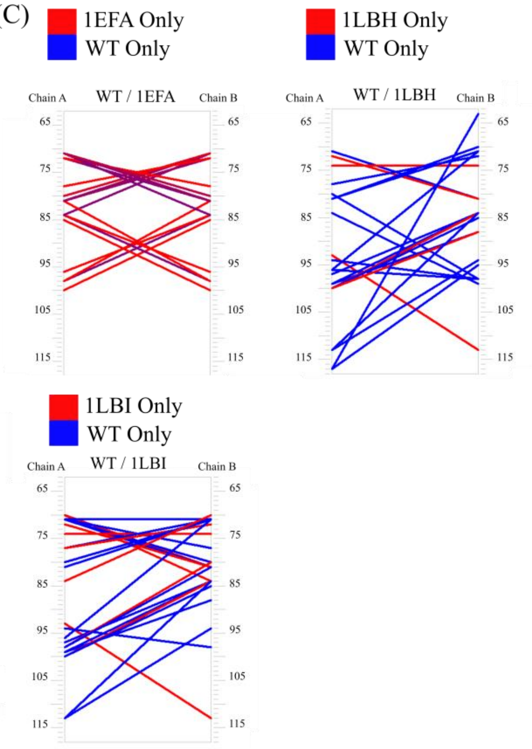

**Figure S2. Structural features of the functional domains in the V52X models.** (A) Overlay of each V52X variant (green, structures extracted from molecular dynamics simulations) with WT LacI (cyan, PDB ID 1efa (Bell and Lewis 2000) using the full structure and chains A and B, individually, of the DNA binding domain along with their associated C-alpha RMSD values. The overall fold of each variant remains largely unchanged

(B) The networks of inter-subunit contacts in the N-subdomains of the V52X variant regulatory domains were determined with Resmap (Swint-Kruse et al. 2001; Swint-Kruse 2004; Swint-Kruse and Brown 2005). Lines indicate inter-subunit contacts; red lines indicate interactions of 3.5 Å or less, whereas blue lines represent contacts between 5 Å and 8 Å; there were no inter-subunit contacts between 3.5 Å and 5 Å for these structures. (C) For comparison, network maps of all interactions 8 Å or less for the crystal structures of DNA/ONPF-bound LacI (pdb 1efa (Bell and Lewis 2000)), IPTG-bound LacI (pdb 1lbh (Lewis et al. 1996)), and apo-LacI (1lbi (Lewis et al. 1996)) are shown. In all network maps, each column of numbers corresponds to amino acid positions in one of the two LacI subunits. In Resmap analyses of the modeled V52X structures, we used all inter-subunit interactions observed in the predominant structural cluster during the final 40 ns of the molecular dynamics trajectory. Each variant manifested 3-5 clusters, with the dominant cluster being the one most frequently sampled.

The features of these modeled structures indicate that most of the interactions were similar to those in the crystal structures. Many of the modeled structures (including WT) appear to have transitioned part-way from the DNA binding conformation to the apo/IPTG-bound conformation, as would be expected since DNA was not included in the simulations. However, since the linker hinge helices did not unfold, the structures more closely resembled the DNA-bound conformation than the apo-conformation (Ha et al. 1989; Kalodimos et al. 2001; Spolar and Record 1994; Spronk et al. 1996; Swint-Kruse et al. 1998; Swint-Kruse et al. 2002; Taraban et al. 2008). These structural analyses support our conclusion that modeling the V52X substitutions did not artificially perturb the structures and that the simulations from which they were extracted provide useful trajectories for DFI/DCI calculations.

The covariance matrices obtained from MD simulations and structures used in this analysis can be accessed through the sbozkan GitHub (https://github.com/SBOZKAN/LacI) and DFI code can be available upon request.

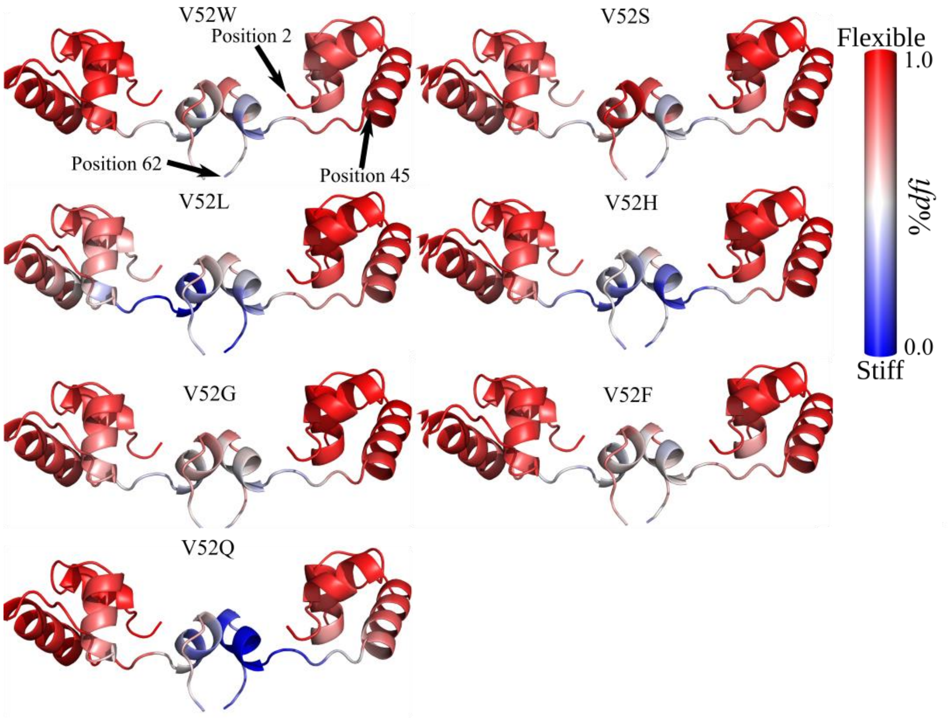


**Figure S3. V52X variants showed altered flexibility.** The DNA binding domains (positions 2-45) and linker regions (positions 46-62) of LacI chains A and B are colored by %DFI. Results for WT and 3 other variants are shown in Figure 4. All results are shown on the WT LacI crystal structure (pdb ID 1efa (Bell and Lewis 2000)). For the seven variants shown here, generally the DNA binding domain remains flexible with subtle changes dispersed throughout. In contrast, the linker regions of these V52X variants stiffen, particularly at the interdomain interface. Plots of %DFI values per position are shown in Figures S4 and S5

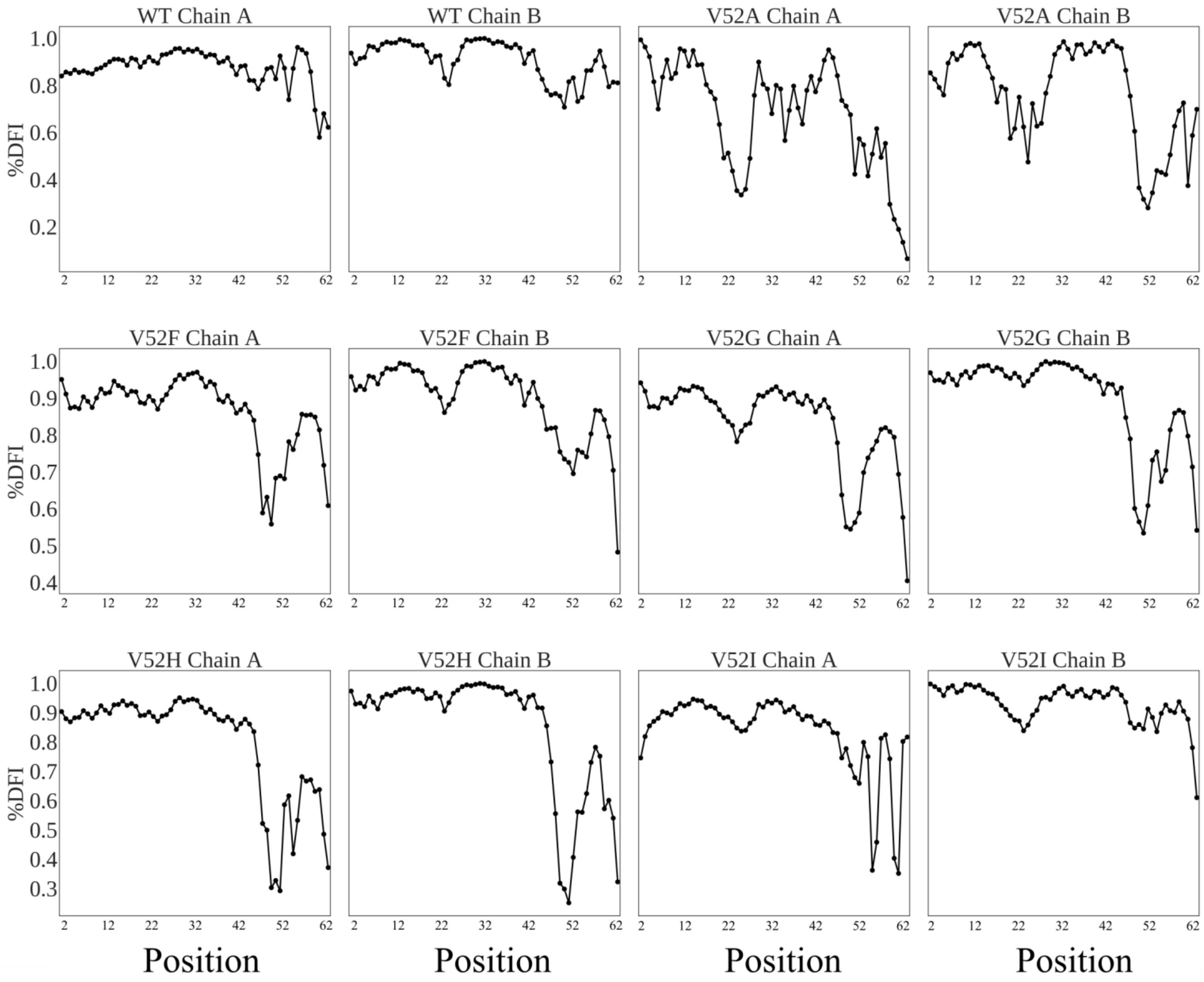


**Figure S4. %DFI scores for all positions in the DNA binding domain and linker region of chains A and B in WT and five V52X variants.** High scores represent high flexibility; the scores for additional V52X variants are in Figure S5. All variants exhibited dramatic flexibility changes near position 52, as expected since position 52 of chains A and B interact with each other, along with other linker positions. In addition, varying degrees of change were observed in the DNA binding domain, particularly around position 22. Notably, the V52A mutation induced dramatic flexibility changes around position 22. Additionally, we find that there is some asymmetry between chains A and B of each variant; although substitutions were made in both chains of the homodimer, the calculations and analysis in the main text was conducted using only one of the chains.

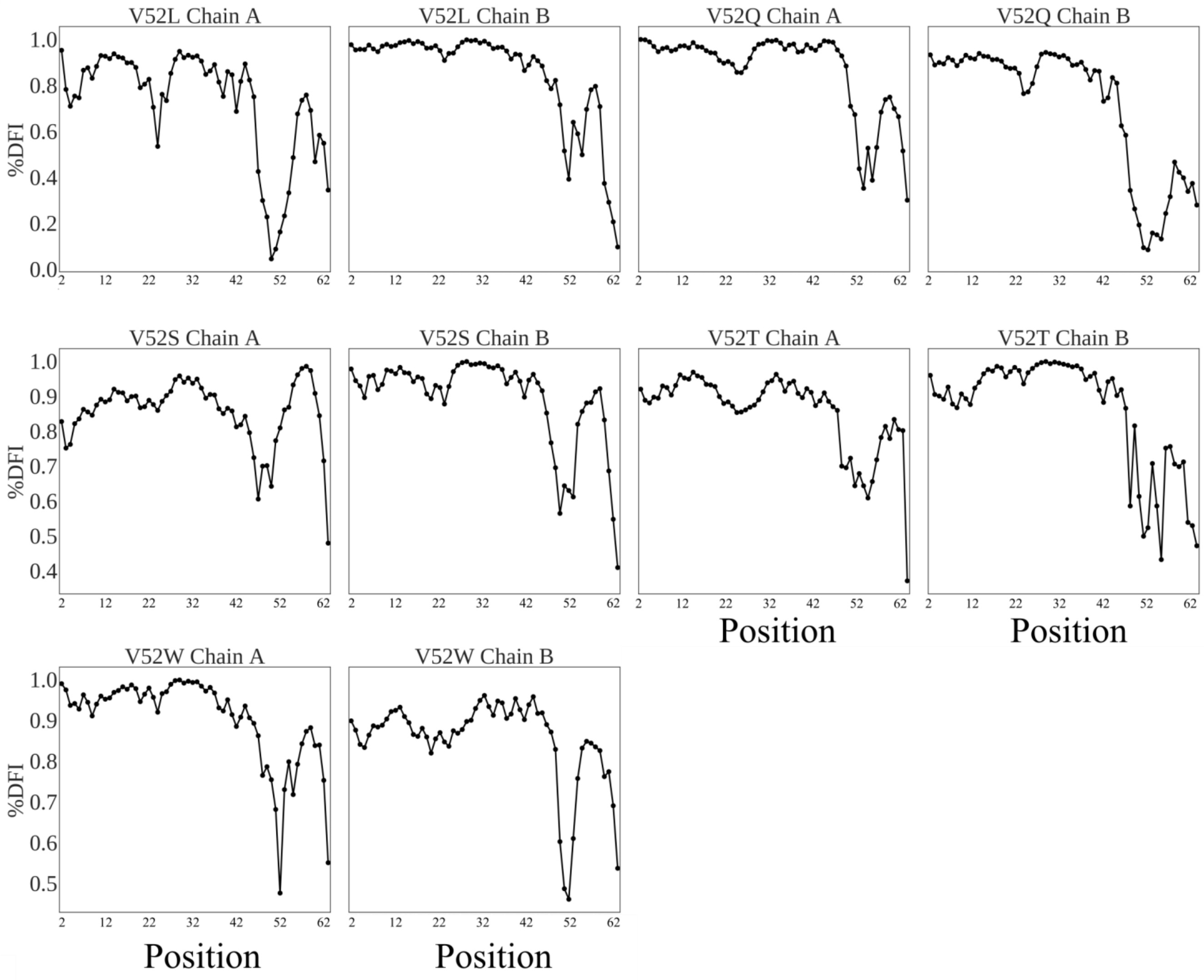


**Figure S5. %DFI scores for all positions in the DNA binding domain and linker region of chains A and B in the remaining five V52X variants.** The features of these plots are described in the legend to Figure S3, which shows the scores for WT and the other five V52X variants.

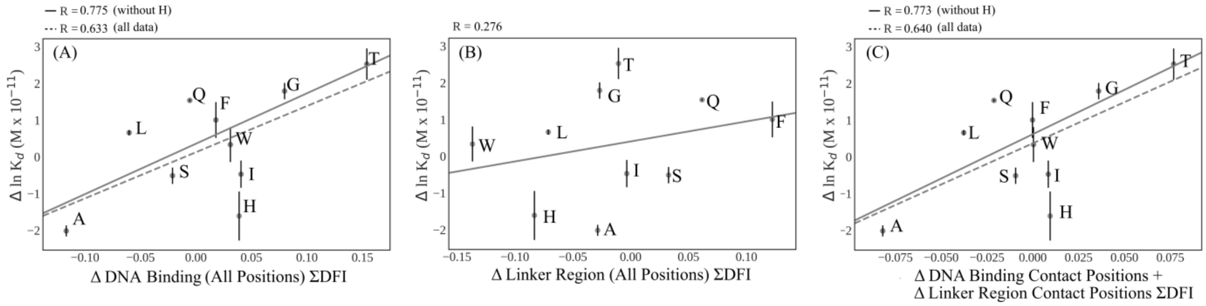


**Figure S6**. **Changes in DNA binding versus changes in flexibility for various subsets of LacI positions.** Regression plots are shown for the change in ΣDFI versus (A) all positions in the DNA binding domains (positions 2-45); (B) all positions in the linker regions (46-62) and (C) all positions in both regions that directly contact DNA (positions 3-7, 14-19, 21, 22, 24, 25, 28-32, 34 in the DNA binding domain and 50, 53, 54, 56, 57, 59 in the linker region). Two additional plots are shown in figure 5 of the main text. For the indicated groups of positions, values for ΔΣDFI are plotted against measured changes in ln $K_d$; changes in both parameters are calculated relative to WT values. (A) The correlation for all DNA binding positions is similar to, and perhaps a little worse than, that of only those positions in direct contact with DNA (Figure 5B). (B) Similar to the linker-DNA contacts (Figure 5C), no significant correlation exists between changes in the DFI scores for all linker positions and ln $K_d$. (C) The correlation for all combined DNA contacts is very similar to that in figure 5B, which indicates that the DNA binding domain contacts dominate the calculation and perhaps dominate DNA binding events.

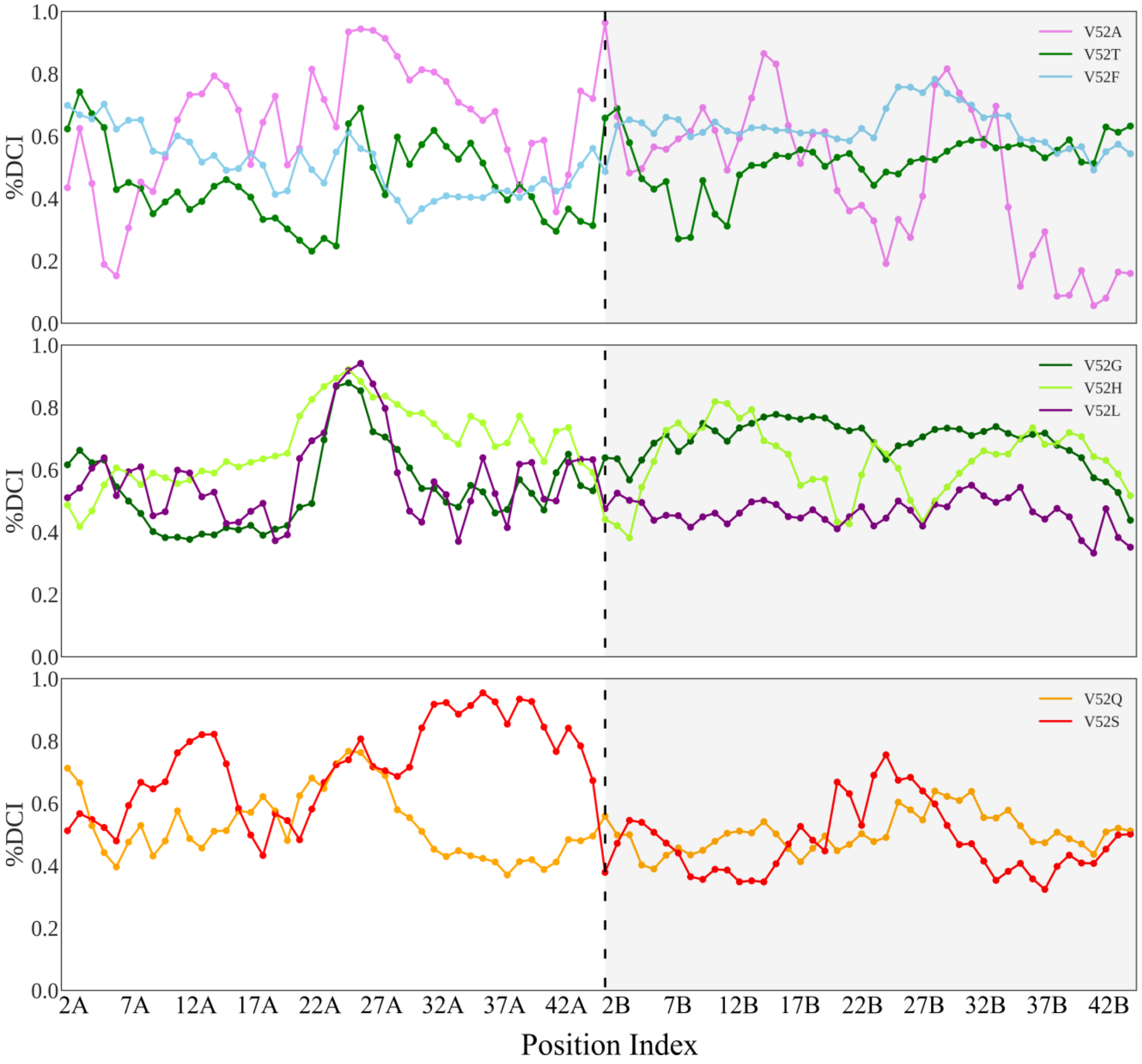


**Figure S7. %DCI scores between position 52 of chain A and all DNA binding domain positions in each of the V52X variants** (those not shown in Figure 7). The "A" and "B" designations on the position numbers and the white/gray shading indicate the two monomers of a dimer. Connecting lines are to aid visual inspection of the data. While each variant exhibited unique coupling behavior, some general trends are evident in many variants, such as high coupling around positions 27A-28A. Additionally, V52S and V52A (which had a tighter DNA binding than WT LacI) have somewhat unique peaks around positions 12-18. V52H, the third variant with tighter DNA binding than WT, also exhibits somewhat higher %DCI values in this region compared to the rest of the variants. Analysis of this data over all V52X variants using principle components from singular value decomposition are shown in figure 6.

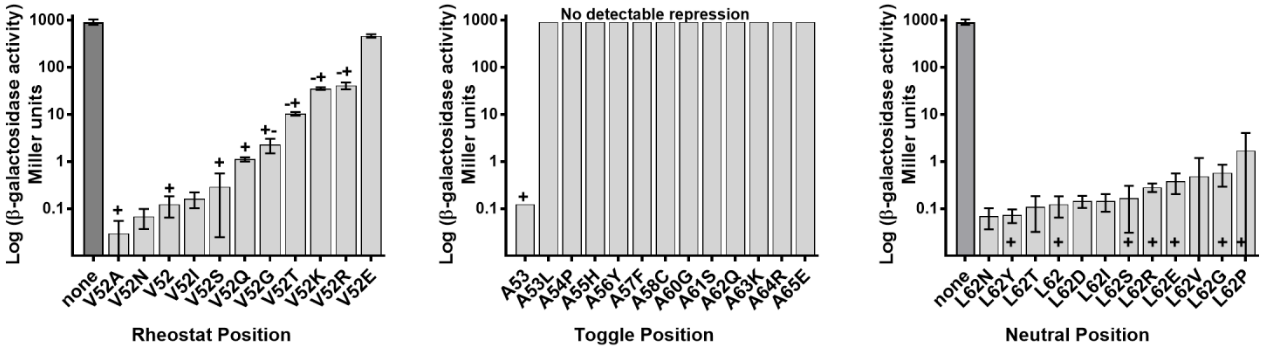


**Figure S8**. **Example positions with rheostat, toggle, and neutral substitution outcomes.** For positions 52 and 62, high-resolution repression data were taken from (Meinhardt et al. 2013) and the “+”, “+/-”, “-/+”, and “-” repression categories are from (Markiewicz et al. 1994; Suckow et al. 1996); these two datasets are further described in Methods. Low bars correspond to strong repression (strong DNA binding affinities). The bar labeled “none” indicates levels of reporter protein activity in the absence of any LacI; as such, it represents a totally “dead” version of LacI. For position 53, data from position 52 were used as benchmarks to recast the low-resolution data onto the high-resolution scale.

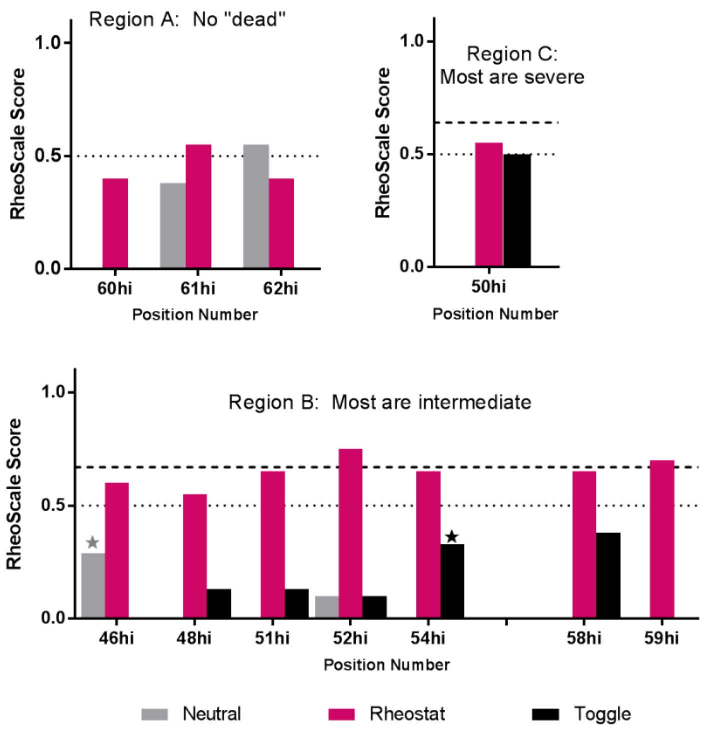


**Figure S9. RheoScale scores calculated from the high resolution functional data of Swint-Kruse et al (Hodges et al. 2018; Meinhardt et al. 2013).** For each position, the experimental outcomes from all available substitutions (*e.g.* supplemental fig. S8 ) were assessed with histogram analyses using the "RheoScale" calculator (Hodges et al. 2018). This calculator describes each position's overall substitution behavior using a set of three scores: "Neutral" scores reflect the fraction of substitutions that have little effect on function. "Rheostat" scores reflect the fraction of the functional range that was accessed by at least one substitution; for the high-resolution dataset, we previously defined rheostat scores ≥0.5 (dotted line) as significant (Hodges et al. 2018). "Toggle" scores reflect the fraction of substitutions that are greatly damaging to function ("dead"); we previously defined toggle scores ≥0.67 (dashed line) as significant (Miller et al. 2017). Two experimental data sets for LacI linker positions are available, each comprising multiple substitutions per linker position; these are further described in Methods. This panel shows RheoScale calculations for the high-resolution repression data that available for nonconserved linker positions; Figure 8 in the main text shows scores calculated from low resolution repression data that are available for all linker positions. The two positions with scores that are over-estimated in the low-resolution data (position 46 has an over-estimated neutral score; position 54 has an over-estimated toggle score) are indicated here with gray and black stars; both exhibit strong rheostat character in the high resolution data, which improves agreement with the %$DCI_{asym}$ analyses shown in Figure 8.

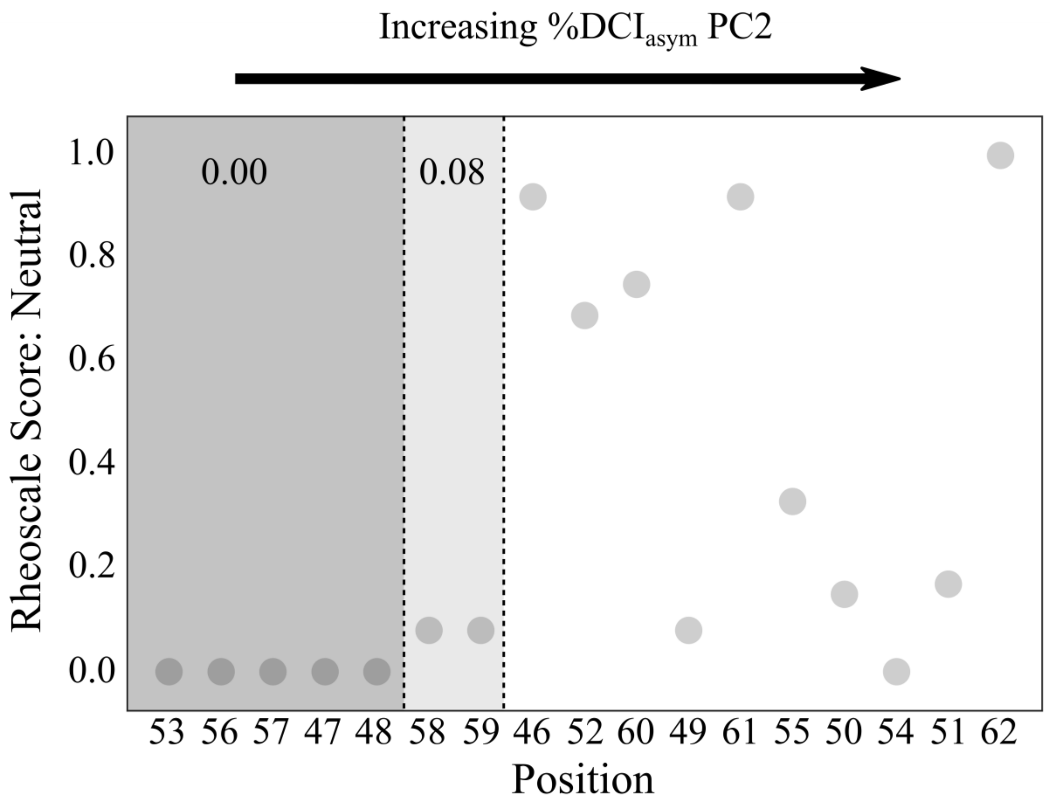


**Figure S10. Linker positions plotted in order of increasing second principle component value from decomposition of DNA-Linker %DCI$_{asym}$.** Principle component 2 can account for much of the discrimination among the types of substitution behavior: Many of the positions with Rheoscale neutral scores of 0.0 (53, 56, 57, 47, 48) exhibited the lowest values in principle component 2. Rheoscale scores were derived using the low resolution repression data (supplementary fig. S8), as in Figure 8. Dashed lines are to aid visual inspection of the data. The combination of principle components 1 and 2 provided a reasonable method of identifying groups of linker region positions when compared to experimental data (see Figure 8).

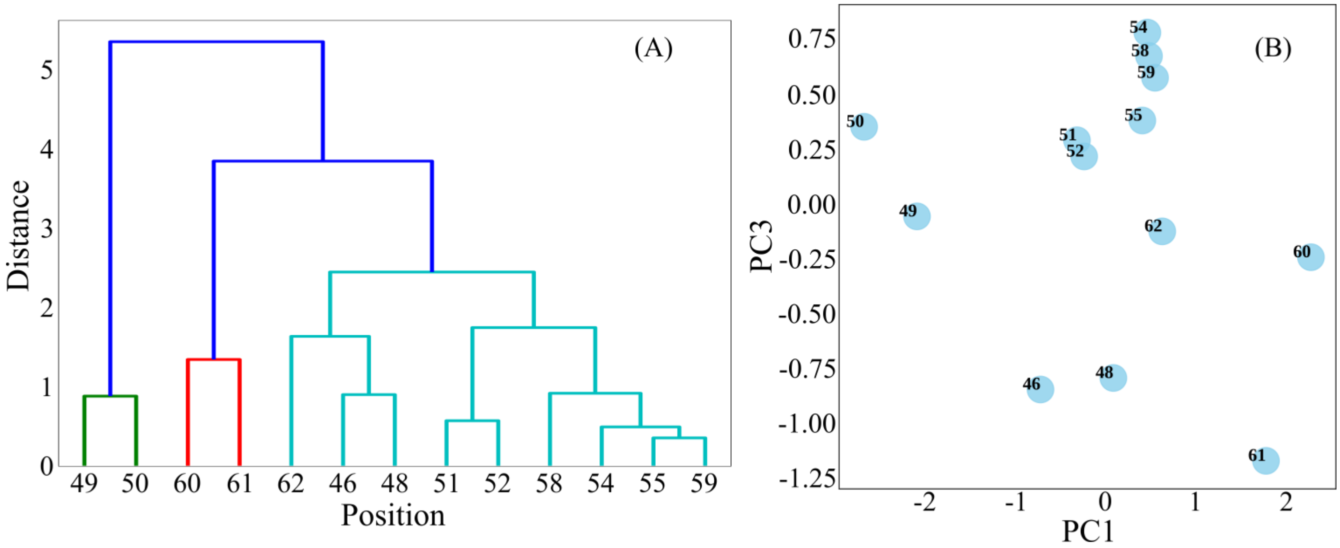


**Figure S11. Dendrogram (A) and PC3 vs PC1 point plot (B) of %DCI$_{asym}$ principle components excluding toggle positions (47, 53, 56, 57)**. (A) Dendrogram of hierarchical clustering (using Ward's method) of PC1, PC3, PC4 and PC5 of non-toggle positions in the linker. PC2 distinguished toggle positions from non-toggles, and further clustering divides the positions into distinct groups, particularly positions 49 and 50 and positions 60 and 61. (B) PC3 vs PC1 of non-toggle positions. Taken together, these plots indicate %DCI$_{asym}$ can group linker positions based on their experimental substitution behavior (see Figure 8).
